# Anti-inflammatory compounds improve spiral ganglion neuron survival after aminoglycoside-induced hair cell loss in rats

**DOI:** 10.1101/2021.12.03.470945

**Authors:** Muhammad T. Rahman, Erin M. Bailey, Benjamin M. Gansemer, Andrew Pieper, J. Robert Manak, Steven H. Green

**Affiliations:** Department of Biology, University of Iowa, Iowa City, IA 52242; Department of Psychiatry, Case Western Reserve University, Cleveland, OH 44106

## Abstract

Spiral ganglion neurons (SGNs) relay auditory information from cochlear hair cells to the central nervous system. After hair cells are destroyed by aminoglycoside antibiotics, SGNs gradually die. However, the reasons for this cochlear neurodegeneration are unclear. We used microarray gene expression profiling to assess transcriptomic changes in the spiral ganglia of kanamycin-deafened and age-matched control rats and found that many of the genes upregulated after deafening are associated with immune/inflammatory responses. In support of this, we observed increased numbers of macrophages in the spiral ganglion of deafened rats. We also found, via CD68 immunoreactivity, an increase in activated macrophages after deafening. An increase in CD68-associated nuclei was observed by postnatal day 23, a time before significant SGN degeneration is observed. Finally, we show that the immunosuppressive drugs dexamethasone and ibuprofen, as well as the NAD salvage pathway activator P7C3, provide at least some neuroprotection post-deafening. Ibuprofen and dexamethasone also decreased the degree of macrophage activation. These results suggest that activated macrophages specifically, and perhaps a more general neuroinflammatory response, are actively contributing to SGN degeneration after hair cell loss.

## Introduction

Spiral ganglion neurons (SGNs) receive input from cochlear hair cells and project from the cochlea to the cochlear nucleus. After hair cell death due to aminoglycoside antibiotics or noise, SGNs gradually die (Green et al., 2008; Spoendlin, 1975). In rats, this death occurs over an approximately three month period (Alam et al., 2007). One possible explanation for SGN death after deafferentation is that they are deprived of neurotrophic factors (NTFs). As we have previously reported (Bailey et al., 2014), while NT-3 expression is greatly reduced after hair cell death, expression of other NTFs, such as CNTF and members of the GDNF family of ligands, persists in the organ of Corti and cochlear nuclei throughout the period during which most SGNs are dying. (We will show here that expression of multiple NTFs also persists in the ganglion itself.) If SGNs continue to be exposed to multiple NTFs after hair cell loss, why do most SGNs ultimately die? Perhaps SGN death is not directly attributable to hair cell loss, but rather to subsequent cochlear events initiated by hair cell loss.

Immune cells, primarily macrophages, are recruited to and activated in the spiral ganglion after noise exposure (Tornabene et al., 2006), diphtheria toxin induced hair cell loss (Kaur et al., 2015), and aminoglycoside exposure (Kaur et al., 2018) in mice. While some studies suggest that macrophages, and fractalkine signaling specifically, are neuroprotective (Kaur et al., 2015), others indicate the presence of proinflammatory signaling, suggesting that immune/inflammatory responses may contribute to SGN degeneration after cochlear trauma (Warchol, 2019; Wood et al., 2017). Immune/inflammatory response activation is known to play a role in other neurodegenerative disorders, including peripheral neuropathies (Royer et al., 2019) and neurodegenerative diseases such as Alzheimer’s and Parkinson’s diseases (Bartels et al., 2020; Subbarayan et al., 2020).

As a first step in determining why SGNs die after hair cell loss, we used microarray gene expression profiling to identify differences in the spiral ganglion between deafened and age-matched control hearing rats. All cells in the ganglion, SGNs, glia, and any immune cells present, were included as it is not only SGNs but glia as well that are affected by hair cell loss (Hurley et al., 2007; Provenzano et al., 2011) and interactions among SGN and non-neuronal cells contribute to SGN survival (Hansen et al., 2001; Stankovic et al., 2004). We profiled gene expression in ganglia from neonatally deafened and control hearing rats at two ages, postnatal day 32 (P32) and P60. P32 is the earliest time at which there is a significant reduction in the number of SGNs, although more than 80% are still alive. By P60, about 50% of the SGNs have died (Alam et al., 2007). Because SGN degeneration after hair cell loss is more severe in the base (Sugawara et al., 2005) and because many genes have been shown to be differentially regulated depending on cochlear region (Flores-Otero et al., 2007), the ganglia used for this assay were hemisected into apical and basal halves in order to make apex-base comparisons of gene expression.

In comparing changes in gene expression between P32 and P60 rats and between hearing and deafened rats, we find that expression of most genes does not change significantly. However, many of the deafness-related changes that do occur are indicative of infiltration of immune system-related cells into the spiral ganglion. We confirmed this with histological data showing an increased presence and activation of immune cells, primarily macrophages, in the spiral ganglion at the time when SGNs are dying, consistent with previous observations (Ladrech et al., 2007; Sato et al., 2010). Immunosupressant treatments reduce SGN death implying that the presence of these immune cells, while possibly related to neuroprotection or clearance of debris, is responsible, at least in part, for SGN death.

## Methods

### Animals, deafening, and verification

All animals used were Sprague-Dawley rats from our breeding colony or from pregnant dams purchased from Envigo. The day of birth was designated as P0. Both male and female rats were used in this study. Rats were housed under a 12 hr light/dark cycle and had free access to food and water. Protocols and procedures for all animal experiments were approved by the University of Iowa Institutional Animal Care and Use Committee.

The microarray experiment was performed in triplicate (i.e. three “biological repeats,” three separate groups of kanamycin treated and littermate control rats). The experimental timeline is shown in Supplemental Figure 1A. Neonatal rats were deafened by daily intraperitoneal injections of kanamycin (400 mg/kg) from P8-16. We have previously shown that, with this deafening protocol, outer hair cells are rapidly lost during the period of kanamycin treatment and inner hair cells are lost more slowly, but are largely absent from all cochlear turns by P18-P19 (Bailey et al., 2014). Supplemental figure 2 confirms loss of hair cells in this study. For two of the three biological replicates, one cochlea from each rat was used as the source of RNA for the microarray analysis while the contralateral cochlea was used for histological verification of hair cell destruction. Hair cells are lost at approximately the same rate in both cochleae from the same rat (Bailey et al., 2014) and we can therefore assume that if hair cells are absent in one cochlea, hair cells are also absent in the contralateral cochlea used for the microarray analysis. In contrast to the untreated controls (Supplementary Figure 2A), the kanamycin-treated rats had no surviving outer hair cells and few, if any, inner hair cells, neither at P32 (Supplementary Figure 2B) nor at P60 (Supplementary Figure 2C). A few rats had some inner hair cells remaining in the apical third of the cochlea. Any rat with >10 such remaining inner hair cells in the apical region of the cochlea were removed from the study. While our criterion for success in eliminating hair cells was histological observation of hair cell loss in one cochlea, nevertheless, to avoid unnecessary histology, we measured auditory brainstem response (ABR) in all rats at P21 (click stimuli, 1-70 kHz stimulus range). Any kanamycin-treated rats showing detectable wave 1 with stimuli 95 db SPL were excluded from the further study. Rats identified as deaf by this ABR criterion indeed lacked inner and outer hair cells in nearly all cases. Rats used for one of the biological replicates were not assessed histologically but any having a detectable ABR wave 1 at 95 dB SPL were excluded from the study.

### Microarray gene expression profiling

For tissue isolation from neonatally deafened rats and hearing control littermates, the animals were heavily anesthetized with a ketamine (40 mg/kg)/xylazine (4 mg/kg) mixture and euthanized by live decapitation at P32 or P60. Temporal bones were rapidly removed and placed in ice-cold phosphate-buffered saline, pH 7.4 (PBS) for dissection. The bony capsule and spiral ligament were dissected away from the cochlea. The remaining tissue was divided into apical and basal portions and the organ of Corti was separated from the spiral ganglion. Immediately following dissection, spiral ganglia were placed into Trizol (Invitrogen), homogenized using a pellet pestle (RPI), flash frozen in liquid nitrogen, and stored at −80°C.

Total RNA was extracted from spiral ganglia using an RNeasy mini kit (Qiagen). On-column DNase I treatment was performed according to the Qiagen protocol. RNA quality was verified using a Bio-Rad Experion RNA analyzer. RNA samples with an RQI of less than 8.5 were not used for further study. RNA was amplified (3-4X) using an Ambion Message Amp II kit. cDNA was synthesized using a SuperScript II Reverse Transcriptase Second Strand kit. The cDNA was then labeled using Cy3-coupled random nonamers (Roche NimbleGen) and hybridized for 20 hours to Roche NimbleGen HD2 rat gene expression arrays. The 12X135K format has 12 independent arrays, each with 26,419 genes and five probes per gene. Arrays were scanned on an Axon GenePix 4200A microarray scanner (Molecular Devices) and Pair files were generated for each array for subsequent analysis. Three independent biological replicates (represented by 4-6 hearing control and 4-6 kanamycin-treated rats) were used to generate the data, each represented by three technical replicates with R^2^ correlations exceeding 0.99 across the replicates.

Lasergene ArrayStar (version 17, DNASTAR) was used for noise subtraction and quantile normalization using the robust multiarray averaging (RMA) method (Irizarry et al., 2003), as well as determination of fold changes. We used a >2-fold change in expression level as the minimal criterion for what we report as significant. Genes presented here changed >2-fold in either the apical or basal half of the spiral ganglion, or in both halves of the ganglion.

### Gene set enrichment analysis and network visualization

Gene set enrichment analysis (GSEA) was used to identify sets of functionally related genes whose expression changes after deafening. Normalized expression values of all genes from the microarray (as generated in ArrayStar, see above) were used as input for to the GSEA software (Mootha et al., 2003; Subramanian et al., 2005) from Broad Institute (version 4.0.3, https://www.gsea-msigdb.org/gsea/index.jsp), which allows for statistical comparison of expression of defined gene sets across conditions. The most up-to-date gene ontology (GO) biological processes gene set (c5.bp.v7.0.symbols.gmt) from the Molecular Signatures Database was used. For this analysis, comparisons between overall hearing and deafened, P32 hearing and P32 deafened, and P60 hearing and P60 deafened were made; differences between apex and base were not analyzed in this study but have been made available (GEO accession ID).

GSEA results were then used as input to Cytoscape (Shannon et al., 2003) and the Enrichment Map plugin (Merico et al., 2010) to generate network visualizations. GO terms (nodes) were included in the network if they had a GSEA p<0.005 or False Discovery Rate Q<0.1. Overlap or similarity with other terms (edges) were kept in the network if the combined Jaccard and Overlap coefficient was ≤0.375. Clusters of terms were kept largely as created by Enrichment Map; however, some manual editing/curation of the network was required to improve visual quality and separation between clusters. Cluster names were modified from names provided by AutoAnnotate (DOI: 10.5281/zenodo.594859).

### Quantitative real-time PCR

For tissue isolation from neonatally deafened rats and hearing control littermates, the animals were heavily anesthetized with ketamine/xylazine as above and euthanized by live decapitation at P32 or P60. Spiral ganglia were dissected and RNA isolated as described above for microarray profiling.

Immediately following isolation, RNA was reverse transcribed to cDNA using SuperScript II Reverse Transcriptase (Invitrogen) using the provided random hexamer primers. qPCR was done using SYBR Green I Master Mix (Roche) on a Roche Light Cycler 480 with forward and reverse oligonucleotide primers purchased from Integrated DNA Technologies (Coralville, IA). The qPCR primer sequences (Supplementary Table 1) were obtained from the Roche Universal Probe Library (http://lifescience.roche.com/shop/CategoryDisplay?catalogId_10001&tab_&identifier_Universal_Probe_Library&langId_-1) and confirmed by sequencing PCR products. Negative controls were reactions lacking reverse transcriptase and are designated –RT. Similarly prepared cDNA from P6 rat brain was used for positive controls. Relative RNA expression levels were determined by normalization to an average of three reference transcripts: S16, Gapdh and PGK1.

### Treatment with immunosuppressive agents and other compounds

Neonatal rats were deafened, as described above, for the gene expression profiling, by daily systemic kanamycin injections from P8-16 and deafness was verified by lack of detectable ABR. All kanamycin-treated rats had no detectable wave 1 at 95 dB SPL and were used for this study.

For administration of dexamethasone, a potent anti-inflammatory steroid, P21 rats were weaned from their mothers and housed in separate cages. At this time, half of the hearing rats and half of the deafened rats began receiving dexamethasone (0.25 mg/kg/day) in their drinking water. All rats received the antibiotic Baytril (23 mg/L) in their water to prevent secondary infection. Sugar (25 g/L) was added to the water to encourage drinking and mask the taste of the dexamethasone and Baytril. Rats were weighed daily and fresh water was provided to maintain the correct dosage of dexamethasone. Rats receiving dexamethasone were also given the nutritional supplement Nutri-Cal (Tomlyn) to promote weight gain. Although dexamethasone-treated rats were small in size, they appeared outwardly healthy and all survived through the end of the experimental treatment.

P7C3A20, an aminopropyl carbazole that activates the NAD salvage pathway (Wang et al., 2014) and has potent proneurogenic and neuroprotective properties (Tesla et al., 2012), and the NSAID ibuprofen were also assessed for ability to prevent SGN death. After weaning at P21 and verification of deafening by ABR, deafened rats received daily intraperitoneal injections of P7C3A20 (20 mg/kg in DMSO/corn oil), ibuprofen (40 mg/kg in DMSO/corn oil), or P7C3A20 and ibuprofen at the same doses from P22-70. Control deafened rats were injected with a similar amount of vehicle (DMSO/corn oil) only.

### Immunohistochemistry

For immunofluorescent labeling of SGNs and immune/inflammatory cells, rats were deeply anesthetized with ketamine/xylazine as above and euthanized by transcardial perfusion of ice-cold phosphate buffered saline (PBS) followed by 4% paraformaldehyde (PFA) in 0.1 M phosphate buffer. Temporal bones were removed and the cochleae isolated and post-fixed in 4% PFA overnight at 4° C. Cochleae were decalcified for 3-7 days in 0.12 M EDTA at 4°C, with daily changes of the EDTA. Following decalcification, the cochleae were cryo-protected through a sucrose gradient (10%, 15%, 20%, 25%, 30%, 30 minutes each), then rotated in degassed O.C.T. Compound (Tissue-Tek) overnight at 4°C. The cochleae were then infiltrated and degassed in O.C.T under gentle vacuum for 1-2 h and then embedded in O.C.T and flash-frozen for cryosectioning. Serial sections of the indicated thicknesses were cut parallel to the midmodiolar plane on a Leica Cryostat CM1850 and collected on Fisher Superfrost slides. Collected sections were stored at −20°C until immunolabeling was performed.

Prior to immunolabeling, slides were warmed to room temperature (∼20-22°C). Cochlear sections were then permeabilized with 0.5% Triton X-100 in PBS for 15 min (6 µm sections) or 25 min (25 µm sections), washed with PBS (3 x 5 min each wash), and immersed in blocking buffer (5% goat serum (Gibco), 2% bovine serum albumin (Sigma, catalog #A2153), and 0.02% sodium azide (Sigma, catalog #S2002) in PBS) for 3-4 h at room temperature. After blocking, sections were incubated in primary antibody in blocking buffer overnight (∼16 h) at 4°C. After primary antibody application, sections were washed 3 x 5 min in PBS, then incubated in blocking buffer containing secondary antibodies (Alexa Fluor conjugates, 1:400, Invitrogen) for 3-4 h at room temperature. Sections were then washed 3 x 5 min in PBS and nuclei stained with Hoechst 3342 (10 µg/ml in PBS, Sigma) for 20 min at room temperature. Slides were then washed and coverslipped with Fluoro-Gel Mounting Medium with Tris Buffer (catalog #17985-10, Electron Microscopy Sciences). Primary antibodies included in this study were: mouse IgG2a-anti-β-III-tubulin (Tuj1, 1:400, catalog #801202, Biolegend), chicken-anti-NF200 (1:1000, catalog #AB5539, Millipore), or mouse IgG1-anti-NeuN (1:100, catalog #MAB377, Chemicon) to label SGNs, rabbit-anti-Iba1 (1:800, catalog #019-19741, Wako) to label macrophages, mouse IgG1-anti-CD68 (ED1, 1:400, catalog #31360, Abcam) to label activated macrophages, and combined rabbit-anti-myosin VI (1:400, catalog #KA-15, Sigma) and rabbit-anti-myosin VIIa (1:400, catalog #25-6790, Proteus) to label hair cells. In some cases, SGNs were labeled with a “neuronal mix”, which included a combined mixture of mouse-anti-NF-H (NF200, 1:400, catalog #N0142; Sigma), mouse-anti-TUJ1 (1:400, catalog #T8660, Sigma), mouse-anti-NF-M (2H3, 1:400, Developmental Studies Hybridoma Bank, University of Iowa), and mouse-anti-calretinin (1:400, cat #MAB1568, Millipore). Type II SGNs were labeled using rabbit-anti-peripherin (1:200, cat #AB1530, Millipore).

### Imaging and quantitation

For histological analysis of immune cells, 25 µm thick sections from P70 cochlea were labeled, as described above, to visualize neurons (anti-β-III-tubulin), macrophages (anti-Iba1), and activated macrophages (anti-CD68). Fluorescently labeled cochlear sections were imaged on a Leica SPE confocal system using a 40x (0.95 NA) objective, 1x digital zoom, and a z axis-spacing of 1 µm. All exposure/gain settings were kept constant throughout the experiment. From one cochlea from each animal, 3-6 sections were labeled for quantitation of immune cells, with at least 50 µm separating each section to ensure that no cells were counted twice. All image analysis was performed in Fiji/ImageJ (NIH, https://imagej.net/Fiji). All cell counts and quantitation of immunofluorescence were done on maximum intensity z-projections of 3D confocal image stacks. First, the outline of Rosenthal’s canal (RC) was traced to measure the cross-sectional area of the ganglion in each image. To determine whether there was infiltration of immune cells in the spiral ganglion after deafening, the number of Iba1-immunoreactive (Iba1+) cells (presumably macrophages) in each image was counted and the density per mm^2^ calculated using the measured cross-sectional area. Only cells with a visible nucleus were counted.

For quantitation of surviving SGNs, 6 µm thick sections from P60 or P70 cochleae were labeled as described above with the neuronal mix or anti-NF200 and anti-NeuN, respectively. Sections were then imaged on a Zeiss Axiovert 200 inverted microscope with a 20X objective. Five to ten sections from a cochlea from each animal were labeled and analyzed for SGNs, with at least 25 µm separating each section to ensure that no cells were counted twice. Fiji/ImageJ was used to trace and measure the area of the RC cross-section and count the number of SGNs. The number of SGNs was divided by the area to derive the SGN density (SGNs/mm^2^) in each cross-section. Only cells that were definitive labeled and had a visibly stained nucleus were counted.

For all histological analyses, cochlear location was defined by half-turn increments proceeding from the base to the apex as previously described (Kopelovich et al., 2013), starting at the basal-most turn (adjacent to the round window) and proceeding apically (Supplemental Figure 1B). Values from each cochlear region for an animal were averaged together so “n” is the total number of rats used in the study and is less than the total number of cochlear sections scored. Images of control and experimental sections were captured in the same session using identical settings to allow quantitative comparison. An automated code generator (written in Python) assigns each image a random 8-digit code number prior to analysis so individuals doing the cell counting and other analyses are blind to the experimental condition represented by the image.

### Quantitative immunofluorescence of CD68

The same images used for immune cell counts were used for this analysis. To assess macrophage activation, quantitation of CD68 immunofluorescence (IF) within Iba1+ cells was performed using a custom-written macro in Fiji/ImageJ, based on NeuronRead, a semi-automated tool for quantification of fluorescence images originally developed for analysis of neuronal images (Dias et al., 2017). For this study, the macro was modified to improve identification of Iba1+ cells, which have a more complex morphology than neuronal cell bodies, and measure pixel intensity values of a selected marker within each Iba1+ cell. All plugins and functions within the macro are provided with Fiji, including MorphoLibJ (Legland et al., 2016). First, using the Iba1 IF channel, background subtraction was performed, followed by thresholding, at which the point the macro stopped to allow user correction of the threshold. After thresholding, a series of morphological filters were applied, followed by morphological reconstruction and particle analysis to identify Iba1+ objects, or cells, and create a region of interest (ROI) for each. The macro again stopped to allow for user correction of identified cells. Once Iba1+ cells were confirmed, morphological measurements of each ROI were taken. The ROIs were then overlaid on the original unedited CD68 channel and the average pixel intensity value (mean gray value) of CD68 IF measured within in each ROI, thus providing a cell-by-cell measurement of CD68 IF. All measurements from each image were normalized to the background fluorescence of the same image. This was done by measuring fluorescence in three acellular areas in each image and averaging the measurements together, followed by subtracting this background measurement from the mean gray value for each ROI. The code for the macro is available as a supplementary file (supplementary file X).

### Statistical analysis

Statistical analyses for immune cell counts, neuron counts, and quantitative immunofluorescence were performed in GraphPad Prism (version 8.1.2). A one-way or two-way ANOVA with Tukey’s or Dunnet’s multiple comparisons or an unpaired t-test were used for parametric data while a Kruskal-Wallis with Dunn’s multiple comparisons or a Mann-Whitney U test were used for nonparametric data. The tests used for specific comparisons are indicated in figure legends. A p<0.05 was considered significant.

## Results

### Gene expression profiling of the spiral ganglion of hearing vs. kanamycin-deafened rats

We used microarray gene expression profiling to assay gene expression in spiral ganglia isolated from P32 and P60 hearing and kanamycin-deafened littermate rats. P32 was chosen because this is the time at which SGN death is first statistically significant, but nearly all SGNs remain alive (Alam et al., 2007) so are a population of live but deafferented neurons. By P60, about half of the SGNs have died, while the remaining SGNs exhibit reduced prosurvival and increased proapoptotic signaling (Alam et al., 2007). This cell population is presumably in the process of dying as a consequence of earlier deafferentation.

The microarray assay results are based on three separate biological replicates for each of the experimental conditions (P32 Hearing, P32 Deaf, P60 Hearing, and P60 Deaf), biological replicate having been analyzed in triplicate (3 technical replicates). We set a threshold for determining which genes were expressed in the ganglion at significant levels. Our threshold for significant expression is based on the signal intensity for genes known not to be expressed in the cochlea (e.g. olfactory receptors), on the signal intensity for genes known to be expressed in the spiral ganglion but at low levels (e.g. neurotrophin receptors), and on negative controls built into the arrays. This threshold was determined to be a normalized signal intensity of 260. We considered a gene to be expressed in the spiral ganglion if it was above threshold in at least one of the four experimental conditions (hearing P32, hearing P60, deafened P32, deafened P60) in the apex and at least one of the four experimental conditions in the base. This criterion yielded 14,138 genes expressed in the spiral ganglion in at least one of the four experimental conditions.

We did note some genes highly expressed in hair cells (e.g., oncomodulin, Ocm; myosin VI, Myo6; myosin VIIA, Myo7a) present in the list of expressed genes. Although these did exceed our threshold for significant expression, they were expressed at low levels relative to most other genes that exceeded threshold. It is possible that their presence indicates some contamination by organ of Corti of the spiral ganglion preparation but it is also quite possible that these genes are expressed at low levels within the ganglion. For example, recent single cell RNA-seq studies of SGNs have shown that some “typical hair cell genes”, such as Myo6 and Myo7a, are expressed in SGNs (Shrestha et al., 2018). As additional support of the latter possibility, we observed that these genes showed no significant change in expression in deafened rats, lacking all hair cells, and that other hair cell markers (e.g., prestin) are not expressed at levels that exceed threshold.

Another factor that must be considered is that changes in gene expression may be taking place in the rats up to age P60 simply because of maturation as the rats age. These would accompany or, possibly, prevent or augment any changes in gene expression due to deafening. To identify maturation-related changes in gene expression and distinguish them from those due to deafening, we compared control hearing P60 rats to P32 rats, compared hearing and deafened rats of the same age, and, to assess changes over time post-deafening, compared P60 deaf to P32 deaf and P32 hearing. Although there are significant regional and maturational differences in gene expression, which we will briefly summarize, the focus of this paper is on deafness-related changes in gene expression up to P60.

Supplementary Figure 1C-F shows correlations among comparisons of hearing vs. deafened comparisons at each timepoint (P32 and P60) and cochlear location (apex and base). For this and all further comparisons, a >2-fold change in expression level was used as a minimum criterion for what we scored as a significant difference in gene expression. In these graphs, reduced correlation (lower R^2^) indicates a greater difference in expression of individual genes between two conditions. R^2^ values are generally high, showing that most genes are expressed at similar levels in the conditions compared. The number of genes having a >2-fold change for each comparison is listed in Table 1. Note that in all cases the number of genes significantly changing in expression is a relatively small fraction of 14,138 total genes expressed.

The smallest number of changes are observed with comparisons made at P32, either in regional differences in gene expression (10 up and 141 down in apex relative to base) or deafness-related changes in gene expression (97 up and 104 down in the apex; and 20 up and 12 down in the base), implying that maturation is not yet complete at P32 and that cell populations that are different have not yet fully diverged in their gene expression profiles. Indeed, the greatest number of differences are those associated with maturational changes in gene expression; for control ganglia from hearing rats there are 118 upregulated and 368 downregulated genes in the apex and 164 upregulated and 45 downregulated genes in the base when comparing P60 to P32. Moreover, at P60, regional differences in expression (117 up and 581 down in apex relative to base) or deafness-related differences in expression (441 up and 162 down in the apex, and 44 up and 37 down in the base) are much greater than at P32 in terms of the number of genes that are significantly different.

### Gene set enrichment analysis of microarray gene expression profiling

To gain a better understanding of the major functional changes in gene expression between hearing vs. deafened rats, between cochlear apex and base, and during maturation P32-P60, gene set enrichment analysis (GSEA) was performed using normalized signal intensities from all expressed genes in all conditions. GSEA results were then used as input to the Enrichment Map plugin to generate network visualizations for different comparisons. The network visualization allows for clustering of GO categories based on similarity; i.e., genes shared between or among categories. This analysis allows for identification of broad functional categories of genes (functional gene clusters) that are enriched in certain conditions compared to others. This analysis was conducted mainly to compare gene expression in the hearing vs. deafened ganglion, both overall (Figure 1A) and at P32 and P60 separately (Figure 2). Other comparisons, such as apex vs. base and maturational-related differences, were conducted rather on a gene-by-gene basis, as the number of genes identified in such comparisons was much smaller than the number identified in hearing vs. deafened comparisons. Below we will summarize the main observations for several comparisons, with a more extensive description of deafening-related changes.

**Figure 1.**
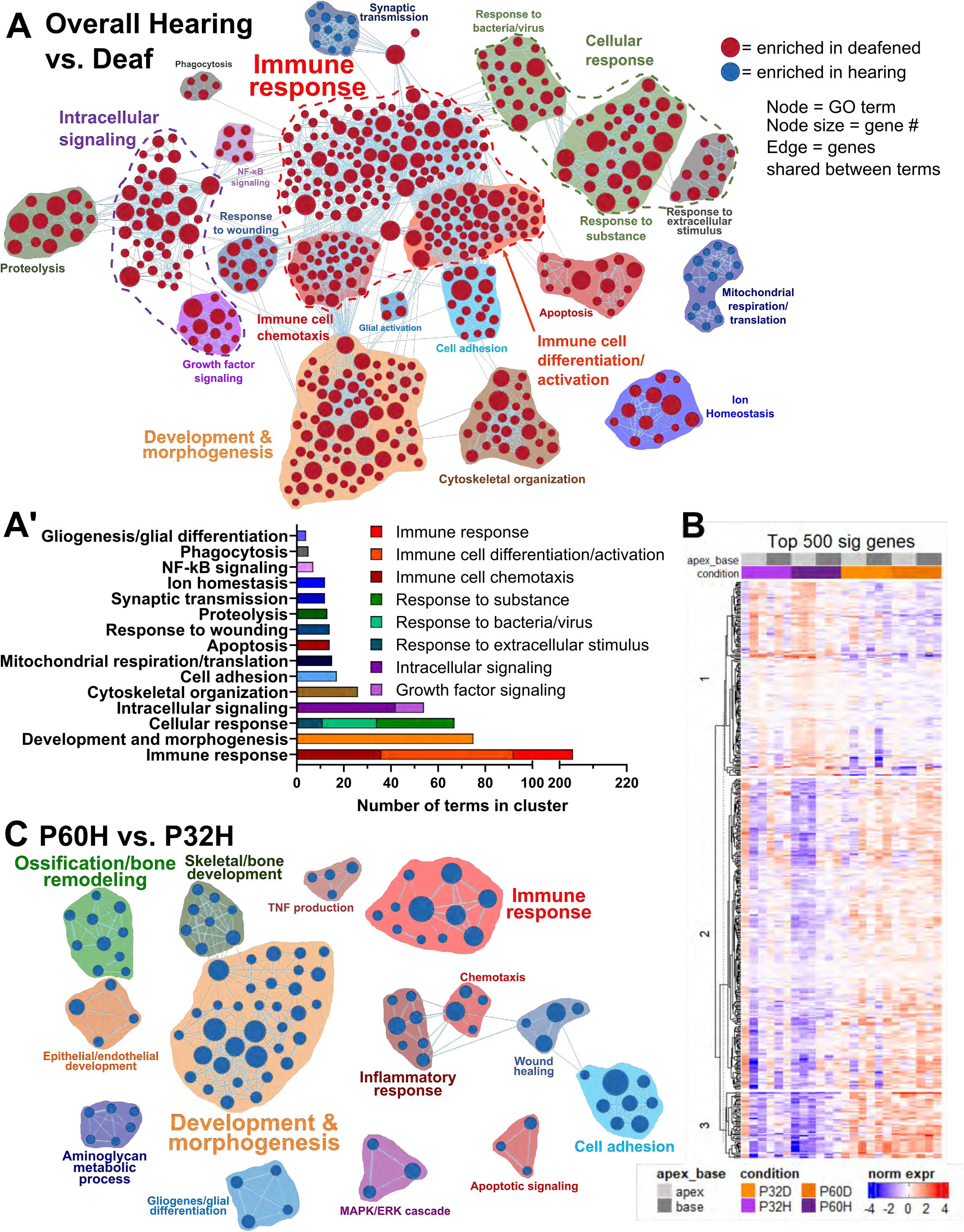
Microarray gene expression profiling reveals differential gene expression in the spiral ganglion of deafened rats. As described in Materials and Methods, normalized signal intensities of all expressed genes in all conditions were used as input to the GSEA software and the results used as input to Cytoscape for network generation. Each node is a Gene Ontology (GO) category, the size of the node is a measure of how many genes are in the category, and edges (lines) indicated that genes are shared between the two connected terms. Nodes were organized into clusters based on similarity (genes shared and category name). **A)** Network showing the overall comparison of hearing vs. deaf, not considering age or cochlear location. Red nodes are categories enriched in deafened while blue nodes are enriched in hearing. **A’)** The number of GO terms in each identified cluster. **B)** Heatmap of normalized signal intensity (further normalized by row) of the 500 genes with largest fold changes (either increase or decrease). **C)** Network showing maturational changes that occur from P32 to P60. Blue nodes indicate that the terms were reduced at P60 relative to P32.

**Figure 2.**
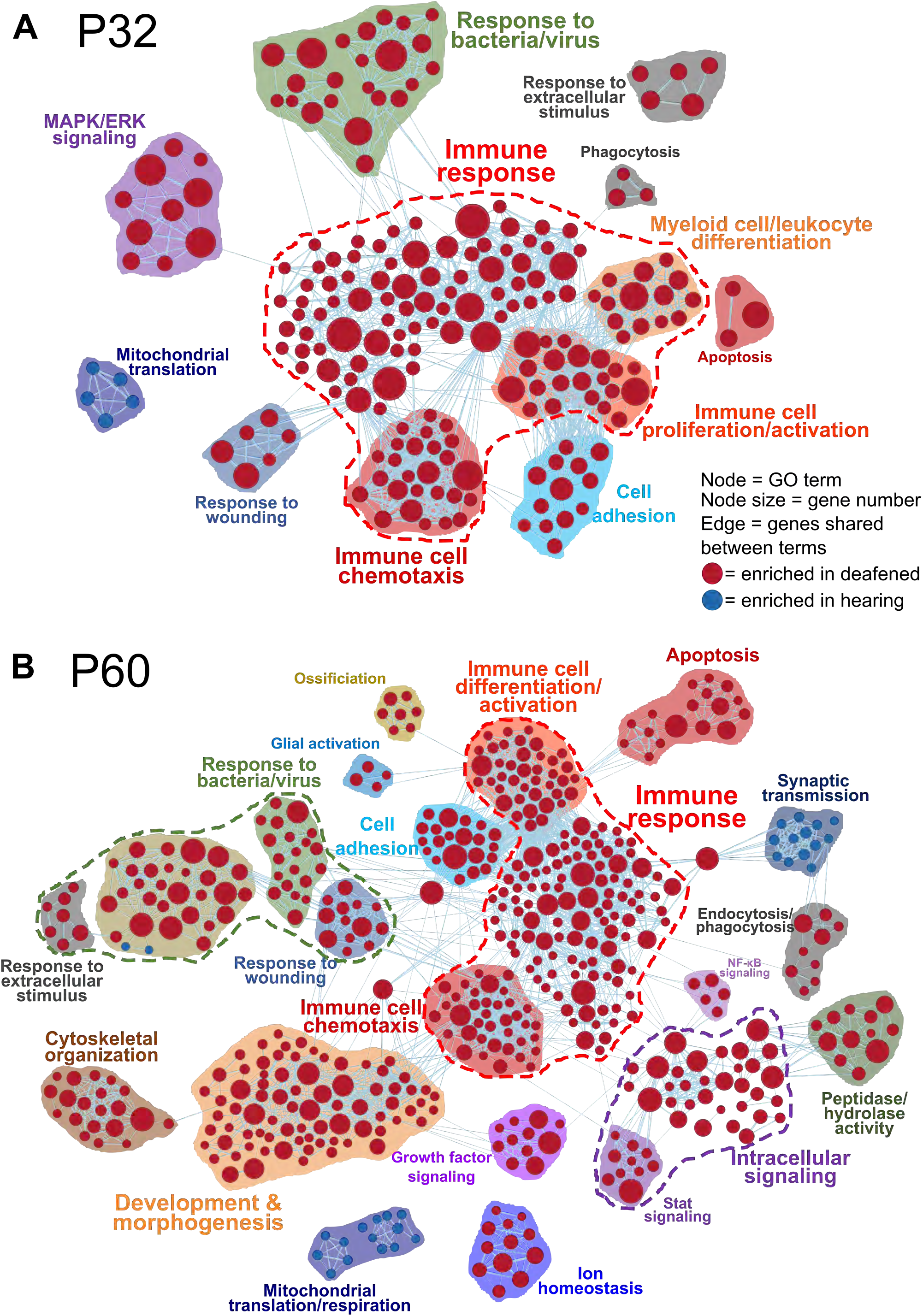
Gene set enrichment analysis (GSEA) of hearing vs. deafened rats at P32 and P60. A) GSEA network showing terms enriched when comparing P32H to P32D. Red nodes indicate enrichment in P32D compared to P32 and blue nodes indicate the opposite. **B)** GSEA network showing terms enriched when comparing P60H to P60D. Note that gene expression changes prominent at P60 are already evident by P32.

#### Apex-Base differences

Divergent patterns in gene expression in the basal half relative to the apical half are evident by P32 (Table 1), a time at which there are 141 genes expressed at lower levels in the apex than in the base and ten at higher levels. The divergence (number of differences) increases by P60 with the same pattern of more genes expressed at lower levels in the apex than expressed at higher levels (117 higher and 581 lower in the apex relative to base). Of potential interest, we note increased expression of p75^NTR^ (Ngfr) and P/Q-type Ca^2+^ channel (Cacna1a, Cav2.1) in the apex relative to base, while claudins and some Ca^2+^ binding proteins are higher in the base relative to the apex. Another category of genes expressed at higher levels in the apex relative to the base are genes encoding presynaptic proteins, including synaptophysin, synapsin 1, synaptojanin 2, synaptotagmin 12, synaptic vesicle glycoproteins 2b and 2c, synaptogyrin 3, and Vglut1, consistent with Flores-Otero et al. (2007). All of these genes encoding presynaptic proteins are expressed at significantly higher levels in the apex relative to the base by P60, but not at P32, indicating that these differences in expression are maturational as well as regional.

#### Maturation-related differences

Genes that change significantly in expression between P32 hearing and P60 hearing are listed in Supplementary Table 2 (upregulated) and Supplementary Table 3 (downregulated). Our data identified 827 genes that change in expression between P32 and P60: 356 upregulated and 471 downregulated. Relatively few of those changes are common to both the apex and base. The numbers of genes that change in expression in the apex, base, or both for up- or downregulation are given in Table 1. As shown in the table, the apex has a much larger number of changes than does the base and most of these are decreases in expression rather than increases. In the base there are fewer changes than in the apex and, in contrast to the apex, most of these changes are increases in expression. It is noteworthy that we observed a much larger number of maturation-related changes in the apex than in the base (618 vs. 341, including genes that change in expression in both apex and base). Cochlear development proceeds in a base to apex direction (Rübsamen et al., 1998) and this may also be the case in this postnatal maturation so some changes we observe in the apex from P32 to P60 may have been initiated earlier in the base.

For genes upregulated at P60 relative to P32, we manually assigned genes to functional categories. With regard to changes common to both the apical and basal halves of the spiral ganglion, 72 genes are upregulated (Supplementary Table 2). In this group, we observe an increase in expression of genes associated with neuronal function. For example, we observe increased expression of several genes associated with ion channel expression and synaptic function. Also present in this group, are genes associated with RNA binding/processing and ubiquitination.

Of 164 genes upregulated only in the base during maturation (Supplementary Table 2), prominent groups include Ca^2+^ binding/ Ca^2+^ signaling, cell proliferation, actin-binding, signal transduction (especially G-protein signaling), immune/inflammation-related, metabolism and biosynthesis (especially relating to lipid), and various transcription factors (especially zinc-finger transcription factors). Also upregulated in the base during maturation are genes encoding ubiquitination-related proteins, additional to those also significantly upregulated in the apex and mentioned above, and 15 plasma membrane proteins, including 10 genes encoding transmembrane transporters. Among the latter is a glial glutamate transporter, which could be related to the high level of synaptic transmission in the cochlear base. We observe a higher expression of glial glutamate transporters in the base than in the apex.

Corresponding with more genes being downregulated, GSEA comparing P32 hearing to P60 hearing identified functional categories/clusters of genes as being enriched in P32 hearing, that is, the genes in these categories are being downregulated from P32 to P60. Overall, 110 GO categories were identified as enriched in P32H (Fig. 1C). Many of the identified clusters contained GO categories of genes associated with developmental processes, including general development and morphogenesis (34 categories), ossification/bone remodeling (10 categories), skeletal/bone development (8 categories), and epithelial/endothelial development (4). These groups contain genes associated with extracellular matrix (ECM) and ECM receptors and proteases, suggesting reduced remodeling ECM in the maturing ganglion. Also present in this gene set were genes associated with cell proliferation, cell adhesion, plasma membrane proteins/transporters, transcription factors, and intracellular signaling (i.e., G-protein and MAP kinase signaling). An especially prominent group of genes downregulated from P32 to P60 contains genes associated with immune (11 categories) and/or inflammatory (7) responses (Fig. 1C). These functional groups are also among the most prominent among genes *upregulated* after deafening. Supplemental table 3 summarizes all genes downregulated from P32 to P60 and their assigned function categories.

#### Deafness-related differences

Our main focus in this report is on gene expression changes due to deafening. Genes that change in expression after deafening are listed in Supplementary Tables 4–7. As shown in Table 1, changes in gene expression are already evident at P32 when SGN death is just beginning. The number of changes is much greater at P60, midway through the SGN death period. At both time points, more changes in gene expression are observed in the apex than in the base. Comparing spiral ganglion gene expression in P32 deafened rats (P32D) to P32 hearing rats (P32H), we find 166 total genes expressed at significantly (>2-fold) higher levels and 132 total genes expressed at significantly lower levels at P32D relative to P32H. These are apparently genes, up- or downregulated, respectively, as a consequence of deafening, even at this early time after deafening. Comparing spiral ganglion gene expression in P60 deafened rats (P60D) to P60 hearing rats (P60H), we find 630 total genes expressed at significantly higher levels and 236 total genes expressed at significantly lower levels in P60D relative to P60H.

Figure 1A shows a GSEA network visualization comparing overall hearing vs. deafened, in which 543 GO terms were identified as enriched, 27 in hearing and 516 in deafened. The GO categories were organized into 20 clusters (2 enriched in hearing, 18 in deafened) based on category similarity. Figure 1B shows how many GO categories were in each of the identified clusters; our focus will be on the largest, which contained GO categories related to the immune system/immune responses. This cluster accounts for 40% (205/516) of the categories enriched after deafening, implicating that the majority of genes that are upregulated after deafening are involved in an immune response.

Figure 2 shows GSEA network visualizations identifying functional gene clusters that are enriched in hearing (downregulated after deafening) or deafened (upregulated after deafening) at P32 and at P60.There were 206 GO terms identified as enriched at P32, five in hearing and 201 in deafened. At P60, 521 GO terms were identified, 29 in hearing and 492 in deafened. The number of terms in each cluster is graphed in Supplemental Figure 3. As relatively fewer downregulated genes were identified, there were only two main functional gene clusters identified as enriched in hearing. Notably, the main cluster identified at P60 contained terms associated with synaptic transmission, mainly presynaptic functions, indicating a decline in neuronal functioning after deafening. In contrast with the number of downregulated genes, a large number of genes are upregulated after deafening (Table 1). As shown in Figure 2, there are many functional clusters of genes identified as enriched after deafening at both P32 and P60. Many of the same clusters are present at P32 and at P60, indicating that the major changes observed at P60 have already begun by P32, even though very few SGNs have died by P32. However, the clusters tend to contain fewer GO terms at P32, most likely because there are fewer differentially expressed genes at P32 compared to P60.

Especially prominent among the categories enriched after deafening are those associated with the immune system. These are divided into several subgroups as indicated in Figure 2 and Supplementary Figure 3. Collectively, these clusters account for 69% (138/201) categories enriched after deafening at P32 and 37% (184/492) at P60. Thus, an increase in transcription of immune response-related genes is the most prominent consequence of deafening for gene expression. We also found that a decrease in transcription of immune response-related genes is evident among the changes in gene expression during maturation in the second postnatal month (Figure 1C). We considered the possibility that the apparent increase in transcription of immune response-related genes at P60 in deafened relative to hearing rat spiral ganglia was simply a consequence of failure in the deafened animals to carry out the normal maturational downregulation of the immune response-related genes rather than purely an increase due to deafening. To identify genes for which the transcriptional increase could not be accounted for by failure of maturational downregulation, we identified those genes for which the level of expression in the spiral ganglia of P60D rats was >1.5X the level at P32H rats. This degree of transcriptional increase at P60 cannot be accounted for simply by a failure of maturational downregulation. We found that in either apical or basal halves of the ganglion, half or more of the upregulated immune response-related genes exhibited an upregulation that could not be accounted for solely by failure of maturational downregulation but, rather, was an upregulation induced specifically by deafening (Table 2). Prominent gene groups identified here include genes encoding complement components (C1 components, C2, C3, C4), chemokines and chemokine receptors (Ccl2, Ccl3, Ccl5, Cx3cr1, Ccr5), and immune cell markers (Aif1, Cd4, Cd68, Ptprc/Cd45).

### qPCR verification

We used real-time qPCR of seven representative immune response genes to verify changes in levels of gene expression obtained from microarray profiling (Figure 3). For qPCR verification, gene transcript expression was assayed and normalized to three reference genes: ribosomal protein S16, Gapdh, and PGK1. Five to eight independent repetitions were performed for each age and region. For each biological repeat we assayed expression in pooled ganglia from 3-6 deafened or 3-6 hearing rats.

**Figure 3.**
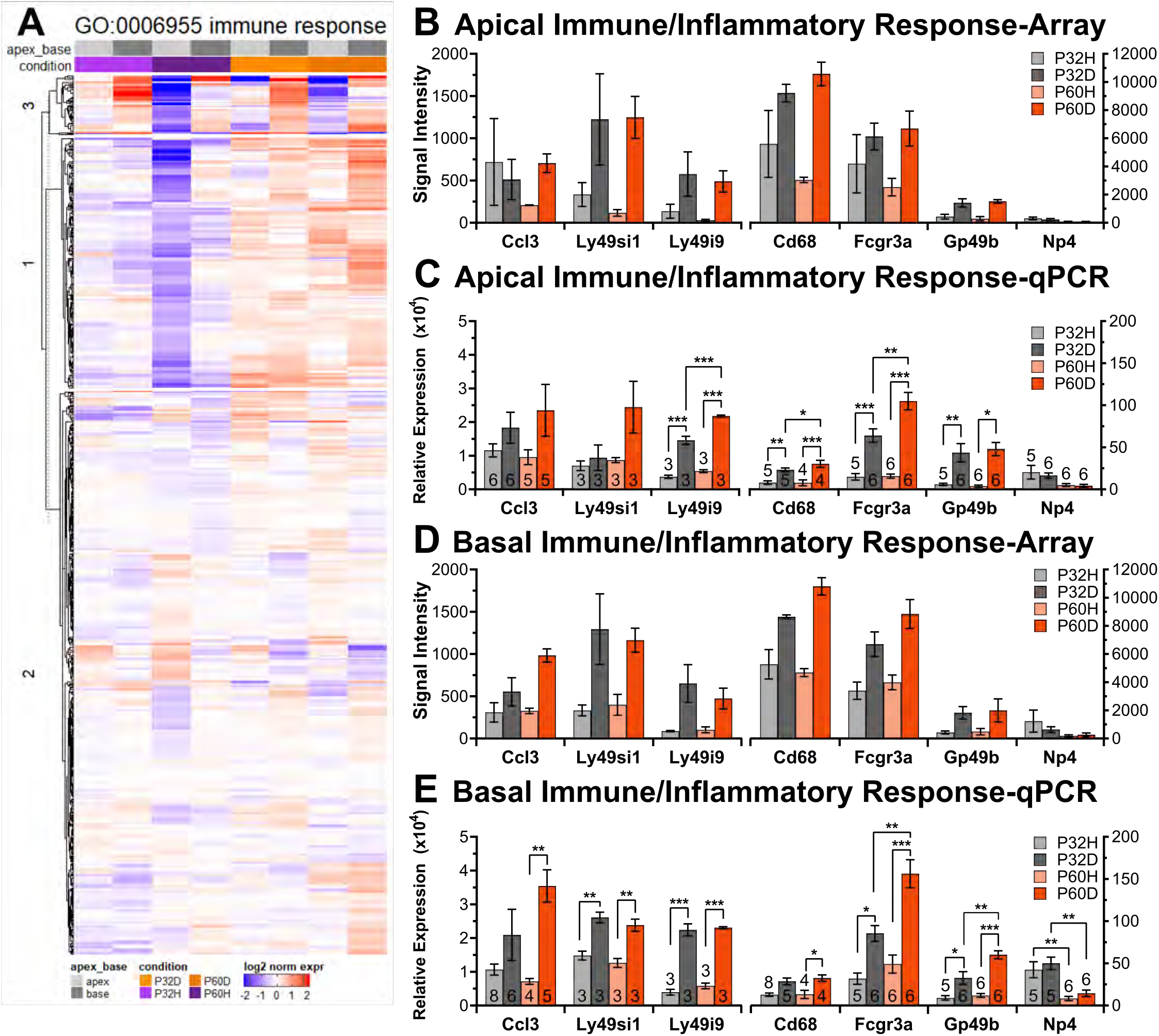
Immune response genes are upregulated after deafening. Heatmap showing normalized expression of genes in the GO category GO:0006955 (immune response) **(A)** indicating that immune/inflammatory response genes are upregulated in the spiral ganglion of neonatally deafened rats. **B-E)** Graphs of microarray signal intensity **(B, D)** and qPCR relative expression **(C, E)** of select immune response-related genes in the apex **(B-C)** or base **(D-E)** across all conditions. n = 3 biological replicates for all microarray data **(B, D)**. One-way ANOVA, *p<0.05, **p<0.01, ***p<0.001; n = # of biological replicates for qPCR data **(C, E)**.

Verification of a change in gene expression was defined as measuring a change in the same direction by both qPCR and microarray. For genes that changed significantly (>2-fold) after deafening or during maturation as assessed by microarray, similar levels or patterns of expression were obtained using qPCR. For example, expression patterns obtained using qPCR showed significantly increased expression of immune response genes after deafening – Cd68, Gp49b, Ly49 receptors – similar to the microarray expression profiles in both the apical and basal spiral ganglia (Figure 3). Changes in genes that were not determined to be significantly regulated using microarray profiling, those that changed less than 2-fold (i.e. Bax, Bcl2, Cntf), were similarly confirmed by qPCR (data not shown).

### Macrophages infiltrate the spiral ganglion after aminoglycoside-induced deafening

Our microarray data show an upregulation of genes expressed in macrophages such as Iba1/Aif1 and CD45/Ptprc after deafening, indicating that macrophage infiltration is occurring. Moreover, our observation of increased expression of genes such as CD68 indicates macrophage activation. It has also been previously shown in mice that macrophages infiltrate the spiral ganglion and other cochlear tissues after noise exposure (Hirose et al., 2005), aminoglycoside exposure (Kaur et al., 2018), and selective ablation of hair cells in the Pou4f3^huDTR^ mouse (Kaur et al., 2015). We sought first to determine whether a similar infiltration occurs in our aminoglycoside-deafened rat model, then determine the effect of neuroprotective compounds or anti-inflammatory drugs on any infiltration. As in the mouse, there is a resident population of macrophages (identified here as Iba1+ cells) present in the spiral ganglion of mature hearing rats (Figure 4). The density of macrophages was similar in all cochlear locations in hearing control animals, an average of 243 cells/mm^2^. Approximately 8 weeks after aminoglycoside exposure there was a significant increase in macrophage density in spiral ganglion in the basal (p<0.05, 2-way ANOVA with Tukey’s post hoc), middle 2 (p<0.001), and apical (p<0.0001) turns. The apparent increase in the middle 1 turn was not significant (p=0.07) (Figure 4). This increase after deafening was more pronounced in the apex, where we observed an average 577 cells/mm^2^, than in the base (382 cells/mm^2^) and middle 1 turn (390 cells/mm^2^). This base to apex gradient in the density of Iba1+ cells in deafened animals was significant (p<0.01), suggesting that there is a more active inflammatory response occurring in the apex at this timepoint.

**Figure 4.**
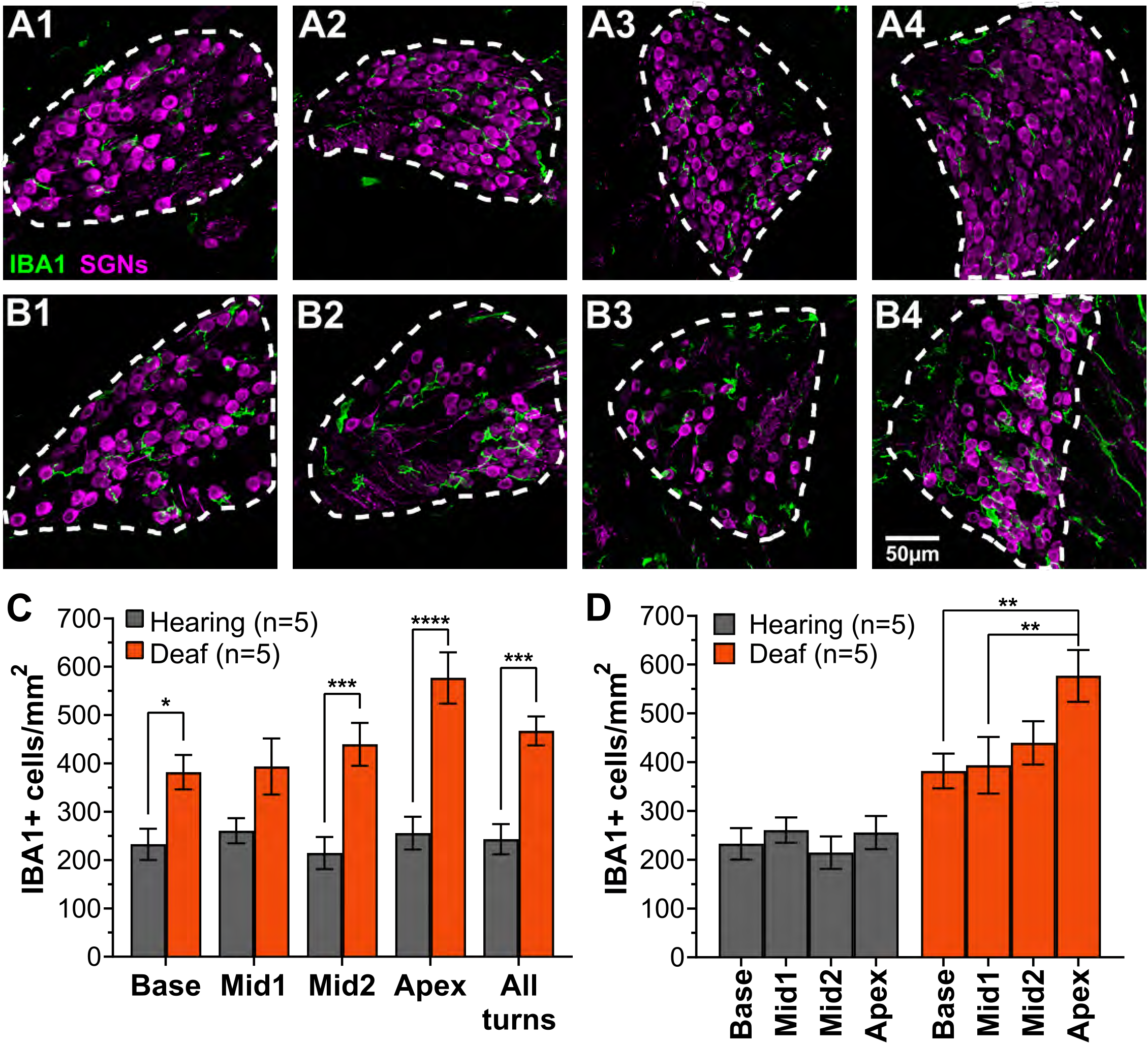
The number of macrophages increases significantly in the spiral ganglion post-deafening. **A**,**B:** Representative images of 25 µm thick cochlear cryosections immunolabeled to show macrophages (Iba1-immunoreactive/Iba1+ cells, green) and neurons (beta3-tubulin – immunoreactive cells, magenta) from control hearing and deafened rats. Rats were deafened P8-P16 and euthanized at P70 as described in Methods. All images are two-dimensional projections of three-dimensional confocal image stacks. Images from P70 hearing **(A)** or deafened **(B)** rats from the basal turn **(1)**, basal portion of middle turn (Mid1**, 2)**, apical portion of middle turn (Mid2, **3)**, and apical turn **(4)**. Dashed lines indicate the outline of Rosenthal’s canal, the border of the spiral ganglion used to determine cross-sectional area in this and all subsequent figures. The number of Iba1+ cells within Rosenthal’s canal in each turn in each section was counted and normalized to the cross-sectional area. **C-D:** Quantitation of the Iba1-immunoreactive cell density (2-way ANOVA, Tukey’s multiple comparisons. *p<0.05, **p<0.01, ***p<0.001, ****p<0.0001; n = # of animals), either comparing hearing to deafened rats at each turn **(C)** or comparing cochlear locations in hearing or deafened cochlea **(D)**.

### Macrophage (Iba1+ cells) activation is increased after deafening

In addition to showing infiltration of macrophages after deafening, we additionally show that there is an increase in the fraction of macrophages that are activated. As noted above, our gene expression data (Figure 3) indicates that genes involved in macrophage activation are upregulated after deafening and that this is evident by P32 when SGNs are just beginning to die. We therefore assessed macrophage activation on a cell-by-cell basis by immunolabeling of activated macrophages. Increased expression of CD68 is associated with phagocytic activity in macrophages (da Silva et al., 1996) so quantifying CD68 IF can be used to determine the degree of macrophage activation. We quantified CD68 IF within individual Iba1+ cells using a macro written in NIH Fiji/ImageJ as described in Methods. Consistent with the microarray results, we found that CD68 IF was indeed significantly increased in Iba1+ cells after deafening (Figure 5). This is evident quantitatively by a rightward shift in the cumulative histogram in deafened animals compared to controls (Figure 5C).

**Figure 5.**
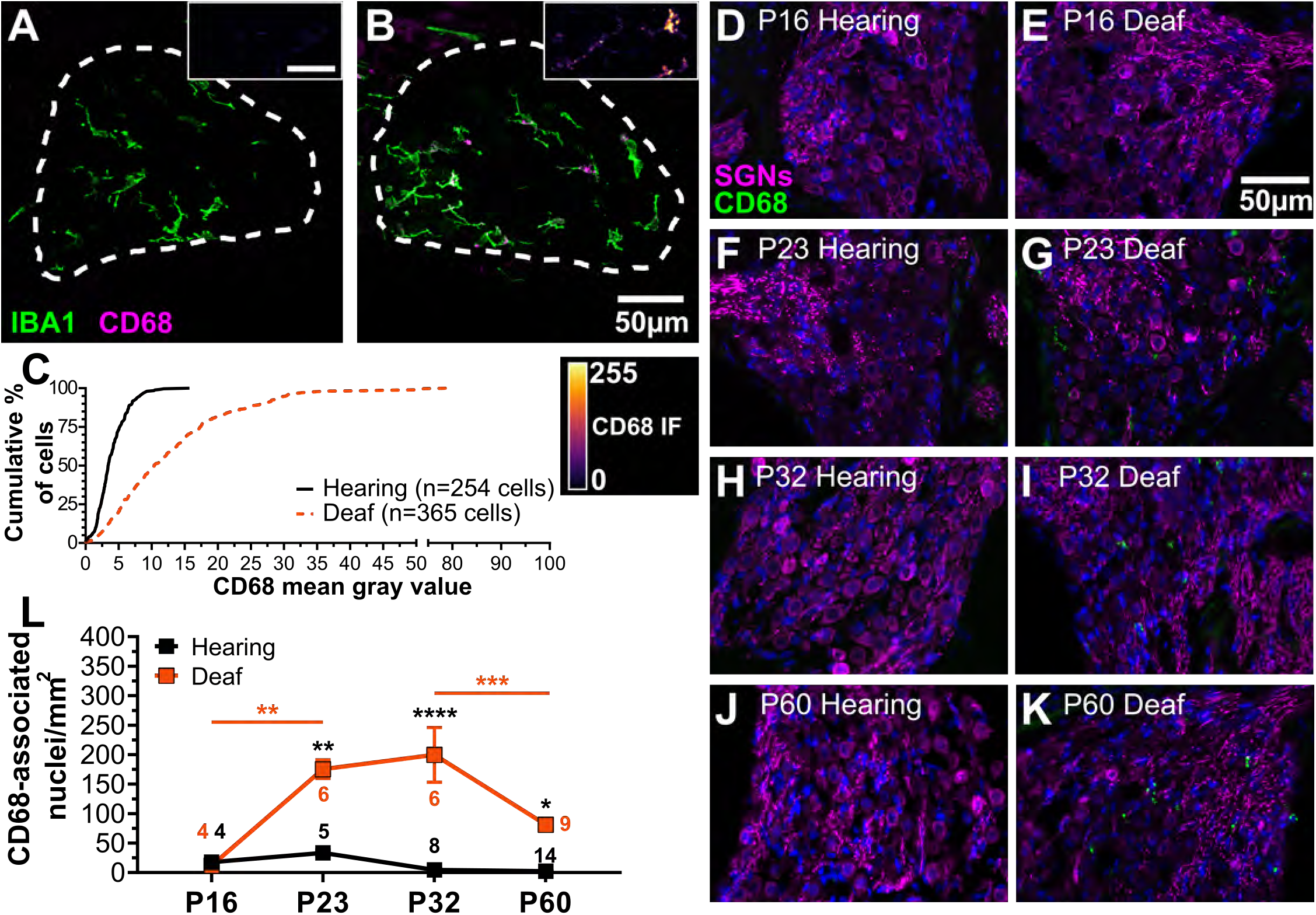
The number of CD68+ cells increases significantly in the spiral ganglion post-deafening. **A,B**: Representative projection images of confocal stacks from the mid 1 turn in spiral ganglia from P70 hearing **(A)** and deafened **(B)** rats showing Iba1 (green) and CD68 (magenta, inferno) labeling. Inset scale = 20µm. Iba1+ cells were identified and the mean gray value of CD68 immunofluorescence in each Iba1+ cell was measured using an ImageJ macro as described in Materials and Methods. A histogram showing the cumulative percentage of Iba1+ cells with a CD68 mean gray value at or above the values on the x-axis is shown in **C**. Note the rightward shift in the curve for the ganglia from deafened rats, indicating a higher frequency of Iba1+ cells with higher CD68 immunoreactivity. **D-K: Timecourse of appearance of Iba1+ cells in the spiral ganglion post-deafening.** Rats, deafened with kanamycin P8-P16, were euthanized at P16 **(D-E)**, P23 **(F-G)**, P32 **(H-I)**, or P60 **(J-K)**. Six µm thick cryosections were labeled with the neuronal mix (magenta) and anti-CD68 (green) as described in Materials and Methods. The number of CD68-associated nuclei/mm^2^ in Rosenthal’s canal was calculated and is shown for each timepoint in **L**. The density of CD68-associated nuclei was compared between hearing and deafened at each timepoint and across the timepoints, within hearing or deafened. 2-way ANOVA, Tukey’s multiple comparisons. *p<0.05, **p<0.01, ***p<0.001, ****p<0.0001; n = # of animals.

Having shown macrophage activation in the spiral ganglion post-deafening, we asked what is its timecourse relative to the timecourse of SGN loss. Note that in rats deafened by kanamycin injections P8-P16, inner hair cells are not entirely lost until approximately P18-P19 (Bailey et al., 2014). CD68+ cells were counted in spiral ganglia from hearing and deafened rats at P16, P23, P32, and P60. The number of CD68-positive macrophages was defined as the number of nuclei adjacent to CD68-positive immunolabeling. Figure 5D-K shows representative examples of immune-labeled sections of Rosenthal’s canal from the mid-apex cochlear region of deafened and hearing rats. Increased CD68-immunoreactivity first appears in deafened ganglia between P16 and P23 (Figure 5D-G) and remains elevated thereafter (Figure 5H-K), consistent with the increase in CD68 gene expression by P32. These quantitative results (CD68-associated nuclei /mm^2^ of Rosenthal’s canal) are shown in Figure 5L Presence of CD68-positive macrophages significantly increases after deafening by P23 and is declining by P60 (Figure 5L), still remaining significantly higher than the hearing control at P60. This pattern of expression is qualitatively similar across all cochlear regions, with the decline in expression at P60 being significant only in the more apical regions of ganglion (Figure 5L). Note that this increase in CD68+ macrophages by P23 occurs *prior* to SGN death, which is not detectable until P28 and not significant until P32 (Alam et al., 2007). This suggests that macrophage activation has some role in the ganglion other than or in addition to phagocytizing debris of dead SGNs and glial cells.

### Dexamethasone improves SGN survival in the base/mid-base after deafening and decreases macrophage activation

To determine whether the immune response might be causing SGN death, we treated rats with dexamethasone, a potent anti-inflammatory and immune suppressant, as described in Methods. We counted surviving neurons in the basal spiral ganglion of hearing control rats, deafened rats, and deafened rats treated with dexamethasone. Representative examples of spiral ganglia sections are shown in Figures 6A-D. Confirming what we have previously shown (Alam et al., 2007), Figure 6E shows a significant decrease in SGN density after deafening in both the base and the basal-most quadrant of the middle region of the cochlea (Mid1). Focusing particularly on these regions, we observed that dexamethasone treatment of deafened rats resulted in significantly higher SGN density than in untreated deafened rats and no significant difference in SGN density between deafened rats treated with dexamethasone and hearing control rats in either the base or mid-base.

**Figure 6.**
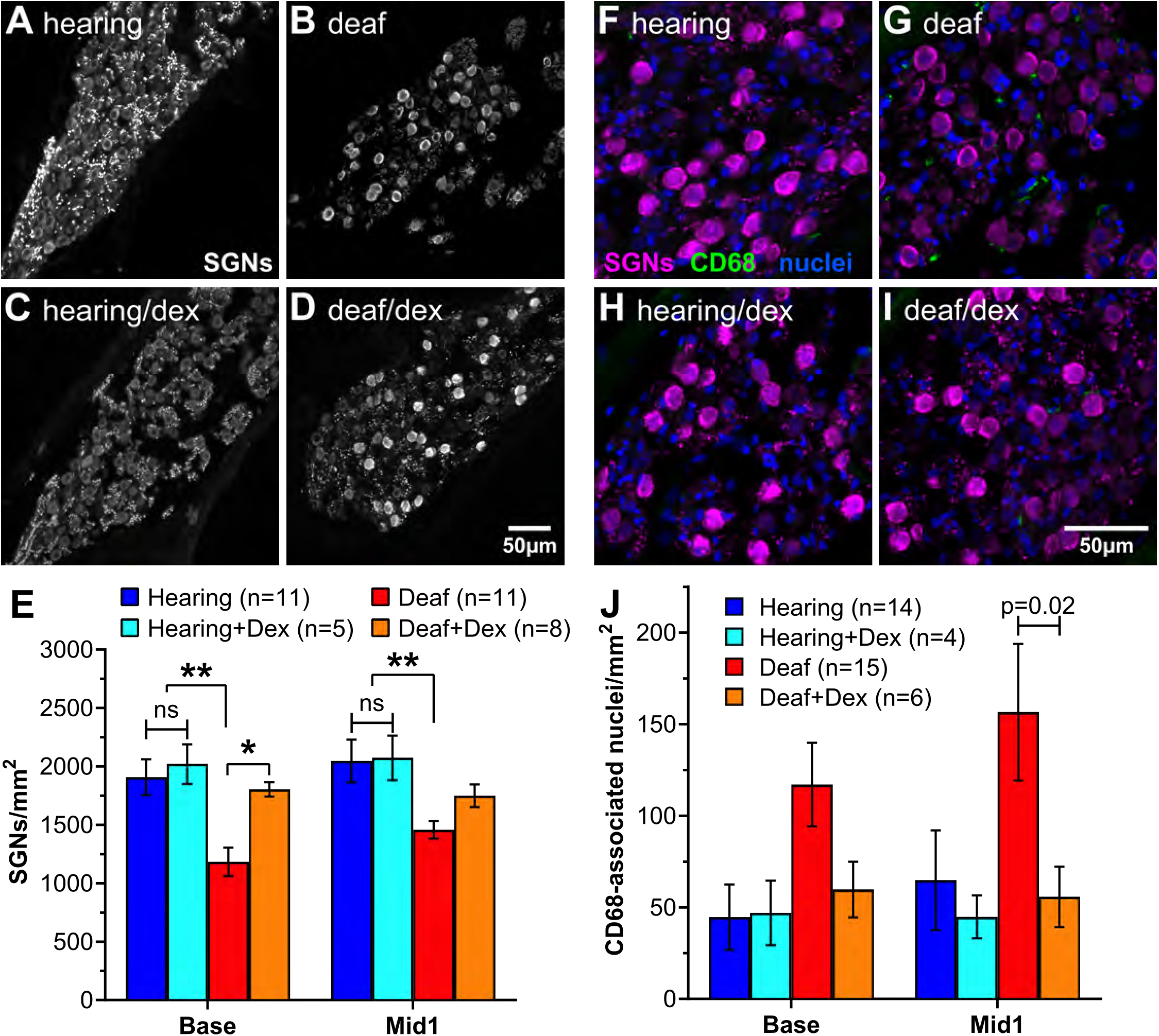
Dexamethasone treatment improves SGN survival in the cochlear base after deafening. Hearing and deafened rats received dexamethasone in their drinking water as described in Materials and Methods. Rats were killed at P60 and cryosections prepared and immunolabeled. **A-D:** Representative images of the basal turn labeled with the neuronal mix to detect SGNs. Scale bar for A-D = 50 µm. **E:** Graph showing quantitation of SGNs/mm^2^ in the spiral ganglion of rats with the indicated experimental conditions. *p<0.05, **p<0.01, 2-way ANOVA, Tukey’s multiple comparisons, n = # of animals. **F-I:** Basal turn labeled to detect SGNs (magenta), nuclei (blue) and CD68 (green) Scale bar for F-I = 50 µm. **J:** Graph showing quantitation of CD68-immunofluorescent cells/mm^2^ in the spiral ganglion of rats with the indicated experimental conditions. Significance determined by Welch’s t-test. n = # of animals.

We next asked whether dexamethasone treatment was in fact able to moderate the post-deafening immune/inflammatory activation. Again, using CD68 immunoreactivity adjacent to macrophage nuclei as a marker for macrophage activation, we counted activated macrophages in control and dexamethasone-treated hearing and deafened rats at P60. Representative examples of spiral ganglia sections are shown in Figures 6F-I, focusing on the basal end of the middle quadrant (Mid1). The counts of CD68-immunoreactive cells are graphed in Figure 6J and show that the post-deafening increase is indeed prevented by dexamethasone treatment. Thus, dexamethasone treatment inhibits the post-deafening immune activation and prevents the concomitant loss of SGNs. These results imply that the post-deafening immune/inflammatory response apparent in the gene expression profiling is causal, at least in part, to the death of the SGNs.

### P7C3 and ibuprofen moderately improve SGN survival after deafening

Our data using dexamethasone implicate immune response activation as a causal factor in SGN death post-deafening. However, long-term systemic treatment with dexamethasone is potentially very detrimental to the health of the subject. We therefore assessed the ability of a potentially safer anti-inflammatory agent, the NSAID ibuprofen, alone or in combination with the neuroprotective compound P7C3 (Pieper et al., 2014), to rescue SGNs post-deafening. Representative images are shown in Figures 7A-F. As shown in Figure 7G, daily ibuprofen treatment reduced SGN loss – i.e., mean SGN density was increased – relative to control deafened untreated rats, significantly in cross-section Mid1. However, ibuprofen was less effective than dexamethasone in reducing SGN loss. P7C3 administration alone appeared to reduce SGN loss but not significantly and less than ibuprofen. Combining ibuprofen and P7C3 improved SGN survival over ibuprofen alone; the increase in SGN density relative to the untreated deafened controls was significant at all cochlear locations. SGN protection was most prominent towards the base in all treatment conditions.

**Figure 7.**
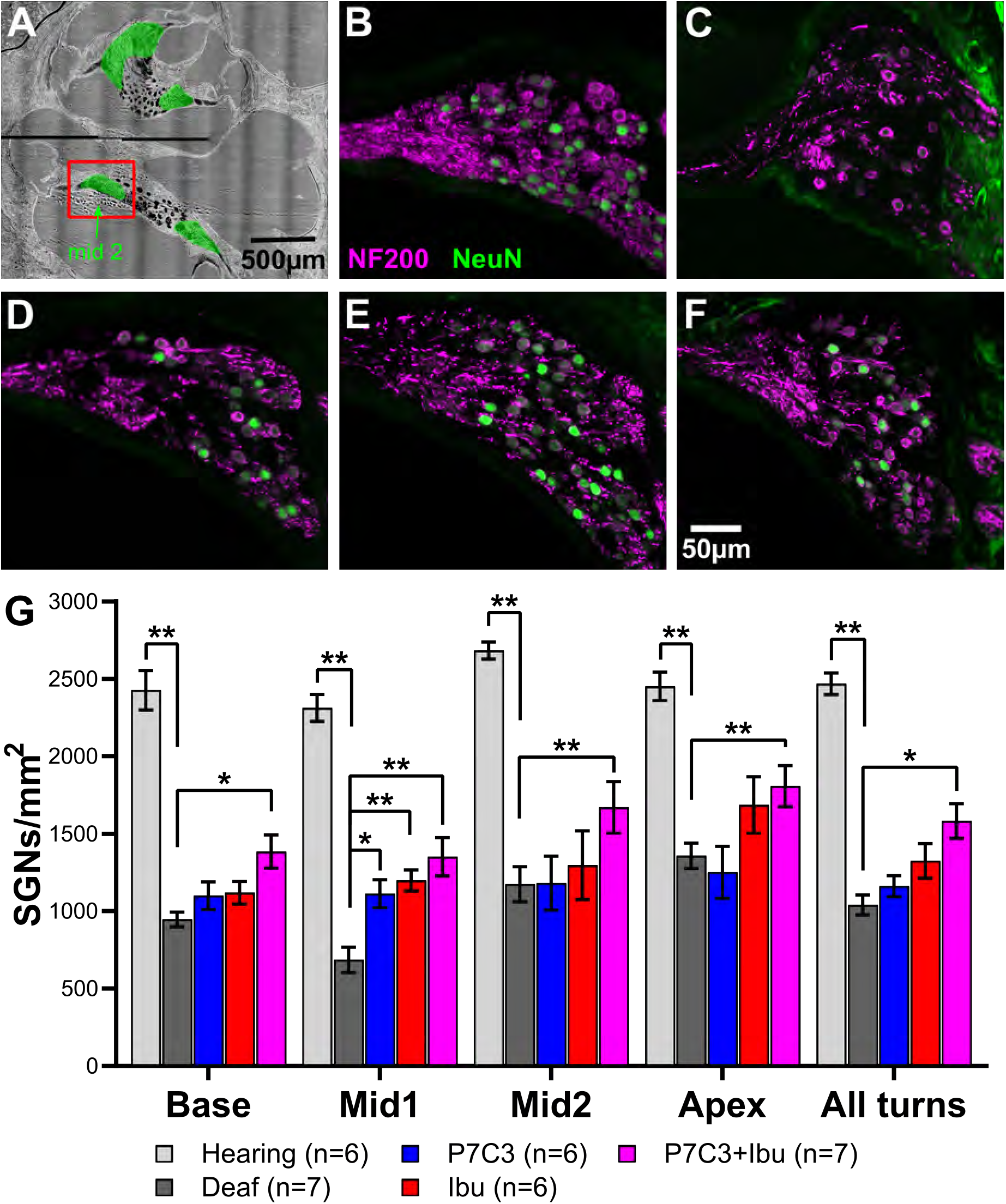
SGN cell density in spiral ganglia from control, ibuprofen- and P7C3-treated P70 rats. Six µm thick cryosections were labeled with anti-NeuN (green) to label SGN nuclei and anti-NF200 (magenta) to label SGN somata and axons. **B-F** are representative images of the middle 2 cross-section (indicated in **A**) of hearing rats **(B)**, or deafened rats treated with vehicle **(C)**, P7C3 **(D)**, ibuprofen **(E)**, or P7C3+ibuprofen **(F)**. Scale bar shown in **F** is for panels **B**-**F**. Quantitation of SGNs/mm^2^ across all conditions and all cochlear turns is shown in **(G)**: 2-way ANOVA, Tukey’s multiple comparisons. *p<0.05, **p<0.01, n = # of animals.

We also investigated the fate of type II SGNs in the ganglion post-deafening. We found that type II SGNs are lost significantly more slowly than type I in the rat, confirming a previous report from (Bichler et al., 1983), so that the fraction of SGNs that is type II rises from ∼5% in control undeafened ears to >10% overall at P90 (Supplementary figure 5). The percentage of SGNs that are type II at P90 in deafened ears was highest in the base (∼17.5%) and declined toward the apex where the percentage of type II SGNs remained near 5%, likely because of the relatively better survival of the type I SGNs in the apex after deafening. In contrast to their effects on type I SGNs, neither ibuprofen nor P7C3, alone or in combination, had any significant effect on the survival of the type II SGNs in any region of the cochlea in deafened rats (Supplementary figure 6).

### P7C3 and ibuprofen do not prevent deafening-induced macrophage infiltration but ibuprofen does reduce macrophage activation

We next asked whether P7C3 and ibuprofen offer neuronal protection after deafening by reducing the immune response. We first assessed macrophage infiltration in deafened rats treated with P7C3, ibuprofen, or a combination of the two drugs. As shown in Figure 8, macrophage density increased significantly in deafened animals from all treatment groups and in all cochlear regions, relative to hearing controls. However, macrophage density was not significantly different among any of the deafened groups, regardless of treatment regimen (Figure 8). Thus, neither P7C3 nor ibuprofen nor both in combination reduced the post-deafening infiltration of macrophages into the spiral ganglion, suggesting that the neuroprotective effects of these compounds are mediated by other means. One possibility is that ibuprofen, an anti-inflammatory agent, suppresses macrophage activation. We assessed this, as above, by quantifying CD68 immunofluorescence in spiral ganglia from hearing and deafened P70 rats. As shown in Figure 9, CD68 IF within Iba1+ cells was significantly increased, relative to hearing controls, in ganglia from deafened rats in all treatment groups. However, ibuprofen partially suppressed the increase: mean CD68 IF within Iba1+ cells in spiral ganglia of ibuprofen-treated deafened rats was significantly lower than in Iba1+ cells in spiral ganglia of deafened rats in the other treatment groups. This result is consistent with the possibility that ibuprofen, an anti-inflammatory compound, reduces SGN death by reducing macrophage activation. In contrast, P7C3 itself did not reduce macrophage activation and, unexpectedly, prevented the reduction caused by ibuprofen. The explanation for this is not obvious but we conjecture that it may be related to P7C3 stimulation of NAD+ synthesis (Wang et al., 2014) that overcomes a deficiency of NAD, required by the immune cells, caused by the significant post-deafening upregulation of CD38, a NAD hydrolase (Aksoy et al., 2006), that we observe in the gene expression data.

**Figure 8.**
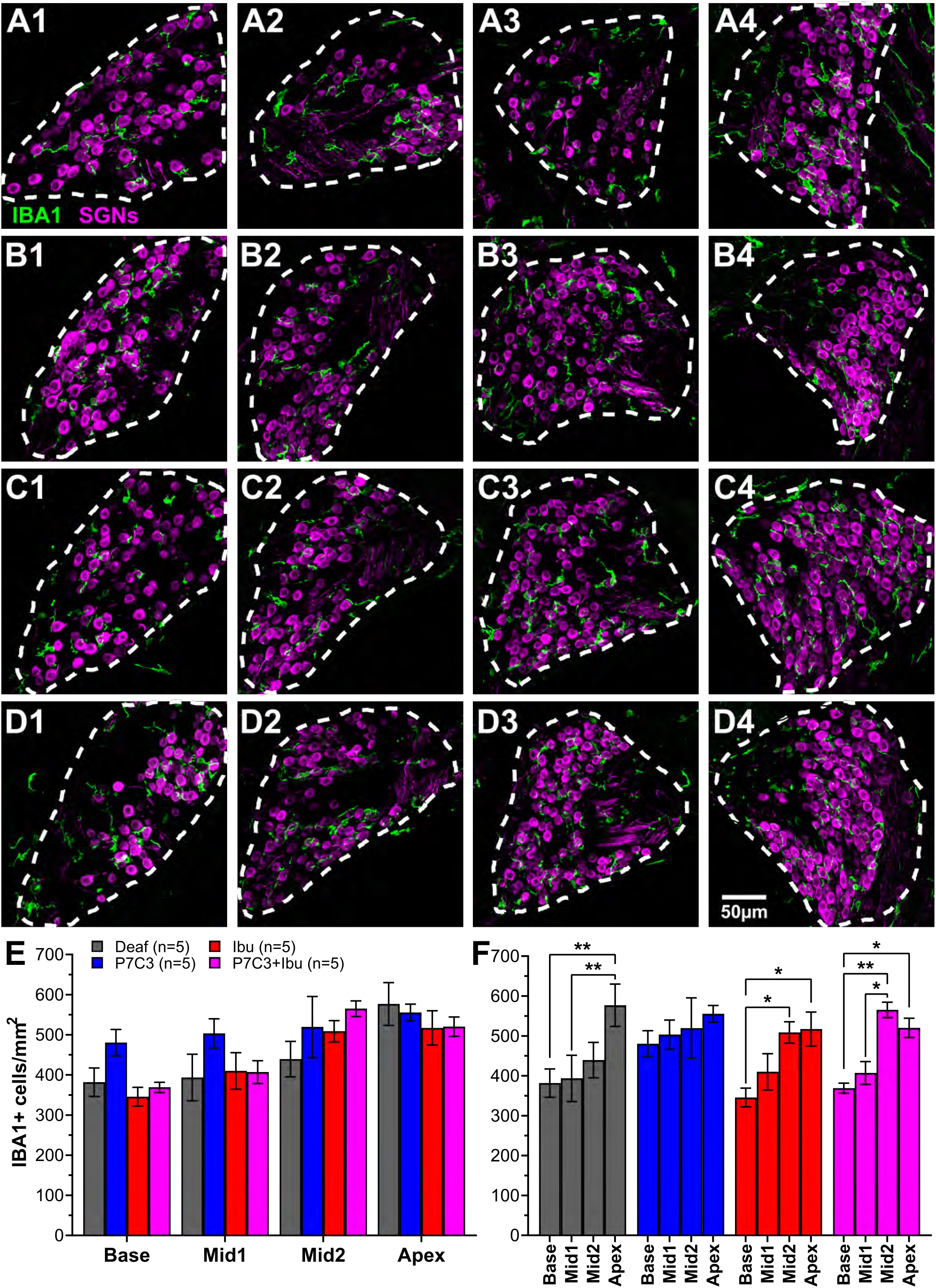
Macrophage number in spiral ganglia of deafened rats treated with ibuprofen and/or P7C3. **A-D:** Representative images of macrophages (Iba1+, green) and SGNs (β-III-tubulin, magenta) from the basal **(1)**, mid1 **(2)**, mid2 **(3)**, and apical **(4)** cross-sections of the spiral ganglion from deafened rats treated with vehicle **(A)**, P7C3 **(B)**, ibuprofen **(C)**, or P7C3+ibuprofen **(D)**. Dashed lines indicate the outline of Rosenthal’s canal/border of the spiral ganglion. All images are projections of confocal stacks. **E**,**F:** Quantitation of the Iba1+ cell density comparing across conditions (treatments) at each turn **(E)** or comparing, within treatment groups, across cochlear location **(F)**. 2-way ANOVA, Tukey’s multiple comparisons. *p<0.05, **p<0.01; n = # of animals.

**Figure 9.**
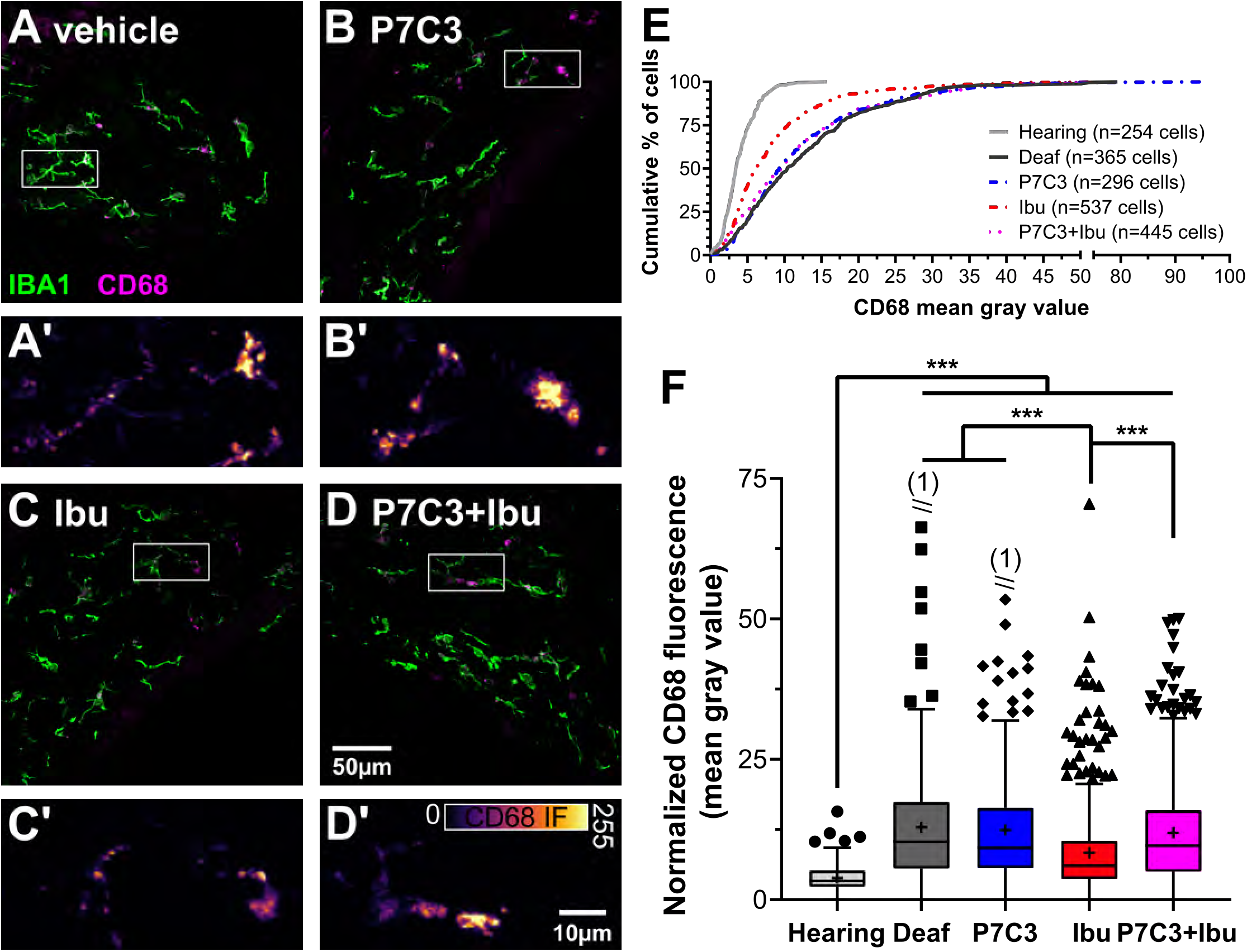
Activated macrophages in spiral ganglia of hearing rats or deafened rats treated with ibuprofen and/or P7C3. **A-D:** Representative images showing Iba1 (magenta) and CD68 (green) immunoreactivity in deafened rats treated with vehicle **(A)**, P7C3 **(B)**, ibuprofen **(C)**, or P7C3+ibuprofen **(D)**. The white rectangles are the areas shown at higher magnification in **A’-D’**. **E:** Histogram showing the cumulative percentage of Iba1+ cells with a mean IF intensity of CD68 at or above the level on the x-axis. **F:** Tukey’s boxplot showing an increase in CD68 IF in all deafened conditions relative to hearing. Ibuprofen significantly reduced CD68 immunoreactivity. Boxes show interquartile range (IQR) with median (line) and mean (+), whiskers extend 1.5 times the IQR. Values above 1.5 times the IQR are plotted individually. Numbers in parentheses indicate the number of values above the axis. One-way ANOVA, Tukey’s multiple comparisons, ****p<0.0001.

## Discussion

SGNs slowly degenerate after hair cell loss caused by aminoglycoside exposure. We provide evidence that immune response activation underlies this neurodegeneration. First, our microarray gene expression analysis indicates that the majority of transcriptomic changes are related to an immune response. Second, consistent with this, histological analyses reveal an increase in macrophage number and activation in the spiral ganglion after deafening. Third, SGN survival was improved in deafened rats treated with immunosuppressants such as dexamethasone or ibuprofen. In addtion, our data on the rate of type II SGN death (Supplemental Figure 5) corroborates previous studies showing that type II SGNs degenerate more slowly than type I SGNs, suggesting that type I SGNs may be preferentially targeted for destruction by the immune system.

Our microarray gene expression profiling encompassed analyses of gene expression not only for changes related to deafening – comparison of hearing vs. deafened ganglia – but also for changes related to postnatal maturation – comparison of P32 vs. P60 ganglia – and tonotopic (apex vs. base) differences. We found that significant differences in gene expression in the cochlear apex vs. base are already evident at P32 but that the number of such differentially-expressed genes is greater at P60 (Table 1), indicating that tonotopic differences in gene expression continue to develop during maturation in the second postnatal month. Many of the genes that are differentially expressed between the apex and base are involved in neuronal function. For example, some Ca^2+^ binding proteins (e.g., Calb1, Caln1, S100a8/9) are expressed at higher levels in the base than the apex, which may reflect a protective mechanism against excitotoxic trauma. Flores-Otero and Davis (Flores-Otero et al., 2011) previously noted that the presynaptic protein synaptophysin is more highly expressed in apical relative to basal SGNs. We can confirm this for P60 rats for synaptophysin (Syp) as well as several other genes encoding presynaptic proteins, e.g., the P/Q-type voltage-gated Ca^2+^ channel alpha subunit (Cacna1a). Nevertheless, some differences exist between our results and those of Adamson et al. (Adamson et al., 2002) for apex-base differences in expression of K_V_3.1 (Kcnc1) and K_V_4.2 (Kcnd2). Such differences in results may be due to the different species, rat vs. mouse, and different ages, P32-P60 vs. P3-P8. We suggest that age-related differences in gene expression extend beyond differences prior to and after hearing onset but gene expression continues to change during maturation until at least two months of age.

The majority of differences in gene expression detected in this study were in the comparison of hearing vs. deafened spiral ganglia at P60. Genes that exhibited difference in expression at P60 between deaf and hearing were further classified as showing a difference because a normal maturational change failed to occur or because expression changed due to deafening in the absence of maturational change. This was done by determining the ratio, not only of P60D to P60H for each gene but also P60D to P32H.

Among the genes up- or down-regulated specifically as a consequence of deafening, we have considered several functional categories. One such category is genes associated with synaptic function. Previous studies in deafened cats have shown synapses made by surviving SGNs in the cochlear nucleus assume a degenerate morphology (Redd et al., 2000). These morphological changes appear to be due to loss of electical activiry in the deafferented SGNs as they are reversible upon electical stimulation of the spiral ganglion (Ryugo et al., 2005), Consistent with these results, we do observe downregulation of expression of genes involved in presynaptic function (e.g., Synj2). Genes involved in axonal conduction are also downregulated post-deafening. The type I voltage-gated Na^+^ channel (the Scn1a gene) is the most highly expressed in SGNs and is significantly downregulated by P60 in deafened ears. Scn8a, but not Scn3a, is also significantly downregulated. Scn7a is abundantly expressed in the spiral ganglion but is reported to be a glial, not a neuronal, Na^+^ channel (Shimizu et al., 2007). With regard to voltage-gated K^+^ channels, many, particularly in the shaker- or Shaw-related subfamilies, are upregulated during maturation between P32 and P60 but this increase is prevented by deafening. Also some voltage-gated K^+^ channels that increase in expression only modestly or not at all during maturation are significantly downregulated after deafening (e.g., Kcna6, Kcnc1, Kcnc2, Kcnc3). Downregulation of these channels is likely to have a significant effect on action potential firing kinetics in the SGNs, with possible implications for the efficacy of cochlear implants. It will be of interest to investigate whether electrical stimulation of the ganglion prevents downregulation of these genes.

A functional category of genes highly relevant to the question of why SGNs die after hair cell loss are genes involved in apoptosis or in neurotrophic factor signaling. We note that apoptosis regulatory genes known to be regulated by neurotrophic stimuli, e.g., Bcl-2, Bax and Bim, do not show significant change in expression post-deafening. This is consistent with our previous conclusion (Bailey et al., 2014) that loss of neurotrophic support is not a principal cause of SGN death after hair cell loss. That conclusion was based on the observation that some neurotrophic factors, e.g., CNTF, remain expressed in the organ of Corti and in the cochlear nucleus even after hair cell loss. Here, we extend those results to the ganglion itself, showing that expression of neurotrophic factors and their receptors in the spiral ganglion does not change significantly after hair cell loss.

Prominent among genes upregulated post-deafening are genes involved with cell proliferation, cell motility, and cell adhesion. With regard to cell proliferation, Provenzano et al. (Provenzano et al., 2011) have shown that spiral ganglion Schwann cells reenter cell cycle after deafening. However, the increased expression of genes involved in cell proliferation, motility, and the region may likely be related to the major category of genes upregulated post-deafening: immune response-related genes. We show that resident quiescent macrophages are present in control undeafened rat spiral ganglia as has been previously shown for avian auditory ganglion (Warchol, 1997) and mouse spiral ganglion (Kaur et al., 2015) but macrophage number (Figure 4) and activation (Figure 5) increases post-deafening. This finding is consistent with previous reports of an increase in macrophages in the cochlea (Hirose et al., 2014) and spiral ganglion (Kaur et al., 2018) of deafened mice. Immune cell infiltration has also been observed after acoustic trauma (Hirose et al., 2005; Kaur et al., 2018), suggesting that an immune response is a typical consequence of cochlear or auditory nerve trauma. This response could account for the upregulation of cell proliferation and cell motility genes, related to possible proliferation of resident macrophages, to inflitration of macrophages, and to migration of activated macrophages within the ganglion.

Indeed, a large fraction of the genes upregulated after deafening are indicative of infiltration of immune cells and an activation of an immune/inflammatory response, particularly innate immune cells. This is similar to observations made after acoustic trauma (Han et al., 2012), in which many genes upregulated after noise exposure sufficient to induce a permanent threshold shift were immune response-related. Inflammation is also associated with age-related hearing loss (Su et al., 2020) in C57BL6 mice. In cd/1 mice, which exhibit accelerated age-related hearing loss, pathways promoting inflammation, such as the TNF-α pathway, are correlated with SGN loss in response to age-related hearing loss (Riva et al., 2007). Chronic inflammation, or “inflammaging”, has also been correlated with decreased hearing performance in humans (Verschuur et al., 2014). Together, these findings indicate that immune response activation and/or inflammation may be a phenomenon common to multiple forms of hearing loss and cochlear damage.

The increase in macrophage number and activation (increased CD68 immunoreactivity) post-deafening occurs concomitantly with neurodegeneration in cochlea, suggesting that macrophages could be directly contributing to SGN death after hair cell loss. This hypothesis is supported by our observation that ibuprofen and dexamethasone, both of which reduce macrophage activation in the cochlea (Figure 9), improve SGN survival in the basal cochlea after deafening. Ibuprofen, a commonly used NSAID, is known to ameliorate neuroinflammation (Lim et al., 2000), exerting its effect by inhibiting cyclooxygenase, thereby reducing the production of pro-inflammatory prostaglandins from arachidonic acid (Bushra et al., 2010). Dexamethasone acts in the same pathway, inhibiting phospholipase A2, which is upstream of arachidonic acid and involved in immune cell activation (Nakano et al., 1990), and also inhibiting cyclooxygenase (Goppelt-Struebe et al., 1989). Both of these drugs could, therefore, be protecting SGNs by directly reducing the neuroinflammation that occurs in the spiral ganglion, with dexamethasone having the stronger effect. However, P7C3 appears to work via a different, more directly neuroprotective mechanism: promoting synthesis of nicotinamide adenine dinucleotide (Wang et al., 2014). The recruitment of two different mechanisms for preventing SGN death accounts for the improved SGN survival in rats treated with both P7C3 and ibuprofen over that in rats treated with either one alone (Figure 6). Neither ibuprofen nor P7C3 affected macrophage infiltration in deafened animals and are, presumably, suppressing the immune response by inhibiting macrophage activation.

Previous studies showed that knockout of the fractalkine receptor CX3CR1 in mice reduced macrophage infiltration into the ganglion resulting in increased SGN death after hair cell loss (Kaur et al., 2015). This suggests that macrophages, and possibly other immune cells, are neuroprotective, in contrast to the results presented here. However, these results serve to highlight the complexity of the immune response, which has multiple functions in conditions of neurodegeneration, including potentially promoting or protecting against neurodegeneration as well as a primary function of clearing cellular debris. Possibly, prevention of fractalkine signaling, which would not necessarily be anti-inflammatory, is disrupts a beneficial effect of macrophages (i.e., clearing cellular debris), although other aspects of the immune/inflammatory response (i.e., antigen presentation and T cell responses) are detrimental. More work is needed to further characterize the deafening-induced immune response and elucidate which immune response mechanisms are neuroprotective vs. neurotoxic.

The results of the gene expression profiling suggest that the immune response includes components other than only macrophages and the innate immune response. Upregulation of genes clearly associated with the adaptive immune response are evident – including complement pathway, MHCII, CD4, CD37, CD72, CD74, CD84, and others – implicating the adaptive immune system. This could account for the specificity of the neurodegeneration that, for example, preferentially targets type I vs. type II SGNs (Supplemental Figure 5). Similarly, the immune response following acoustic trauma includes both innate and adaptive immune components (Rai et al., 2020). Thus, further characterization of the immune response is necessary to fully identify all mechanisms involved. This may be of importance in developing therapeutics to prevent SGN degeneration post-deafening. SGN degeneration after hair cell loss is thought to adversely affect cochlear implant function (Heshmat et al., 2020). Therefore, preventing SGN degeneration may improve the efficacy of cochlear implants.

## Supplemental figure legends

**Supplemental Figure 1.**
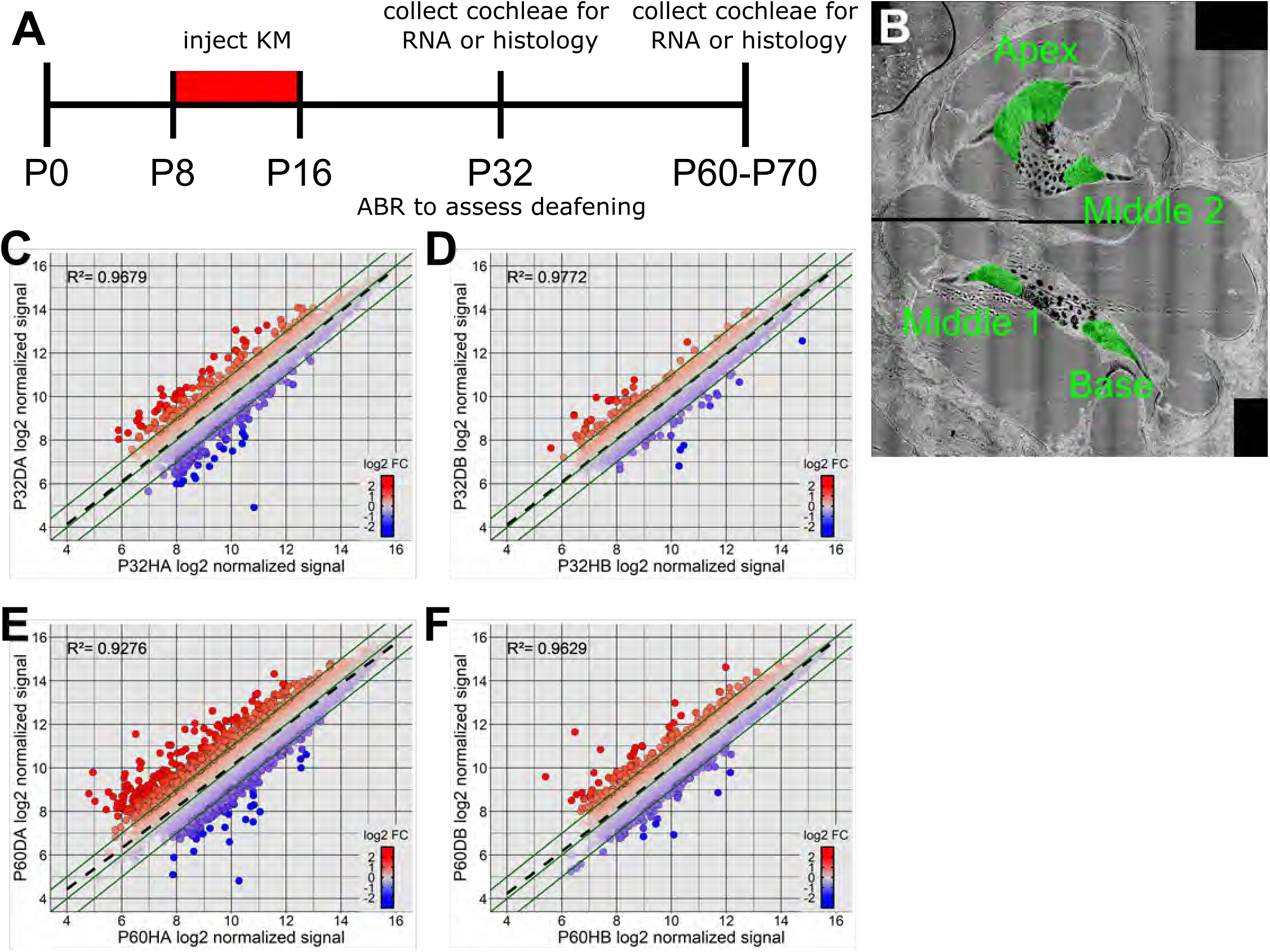
Experimental timeline for the microarray and histological studies. **A:** Rats were injected daily with kanamycin from P8-P16. At P32, ABRs were performed to assess deafening in the injected animals. Additionally, some animals were killed at P32 to collect RNA for microarray analysis. For additional microarray analysis, hearing and deafened rats were killed at P60. For histological studies, hearing and deafened rats were killed at P70. **B:** Representative image of a whole midmodiolar section from a P70 rat cochlea. The different turns and their designations are highlighted in green. **C-F:** Correlation plots comparing normalized signal intensities of all expressed genes between P32 hearing and P32 deaf apex **(C)**, P32 hearing and P32 deaf base **(D)**, P60 hearing and P60 deaf apex **(E)**, and P60 hearing and P60 deaf base **(F)**. Note that in all comparisons the correlation between hearing and deaf is high (R^2^>0.9), indicating relatively few changes in gene expression between the hearing and deafened ganglion. KM = kanamycin; ABR = auditory brainstem response

**Supplemental Figure 2.**
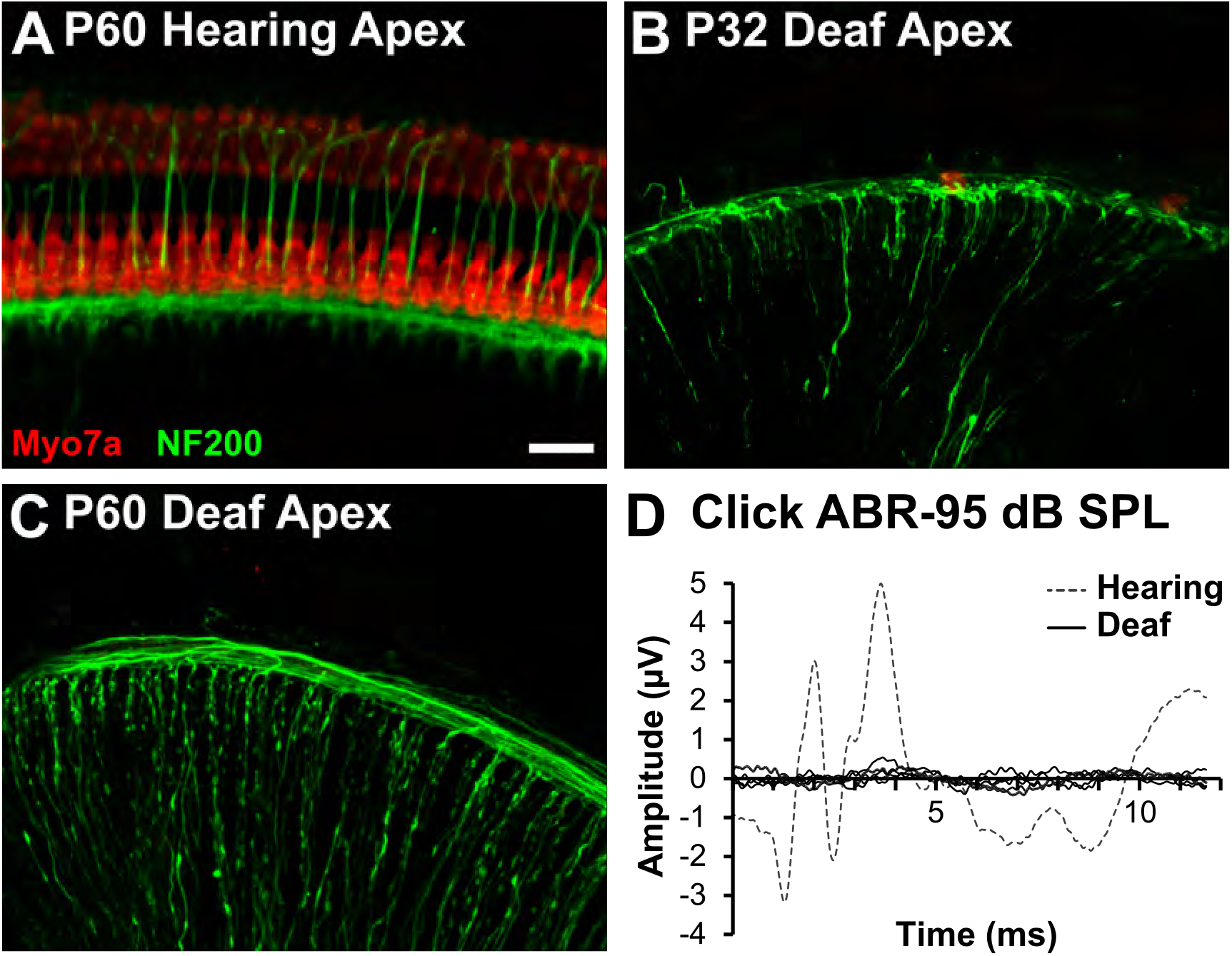
Confirmation of hair cell loss and lack of an ABR in kanamycin-deafened rats. Rats were injected with kanamycin from P8 to P16 as described in Materials and Methods. One cochlea from in each animal in a subset of rats at P32 or P60 was used for histological verification of hair cell loss. Rats were killed and the cochlea removed and prepared for wholemount dissection along with labeling for hair cells (anti-Myo7a, red) and neurons (anti-NF200, green). A normal complement of hair cells is observed in the P60 hearing apex **(A)**, while at P32 deaf **(B)** and P60 deaf **(C)** no hair cells remain. Additionally, ABRs to click stimuli were measured at P32. The representative example in **(D)** shows no detectable ABR to 95 dB-SPL clicks in deafened animals. ABR = auditory brainstem response

**Supplemental Figure 3.**
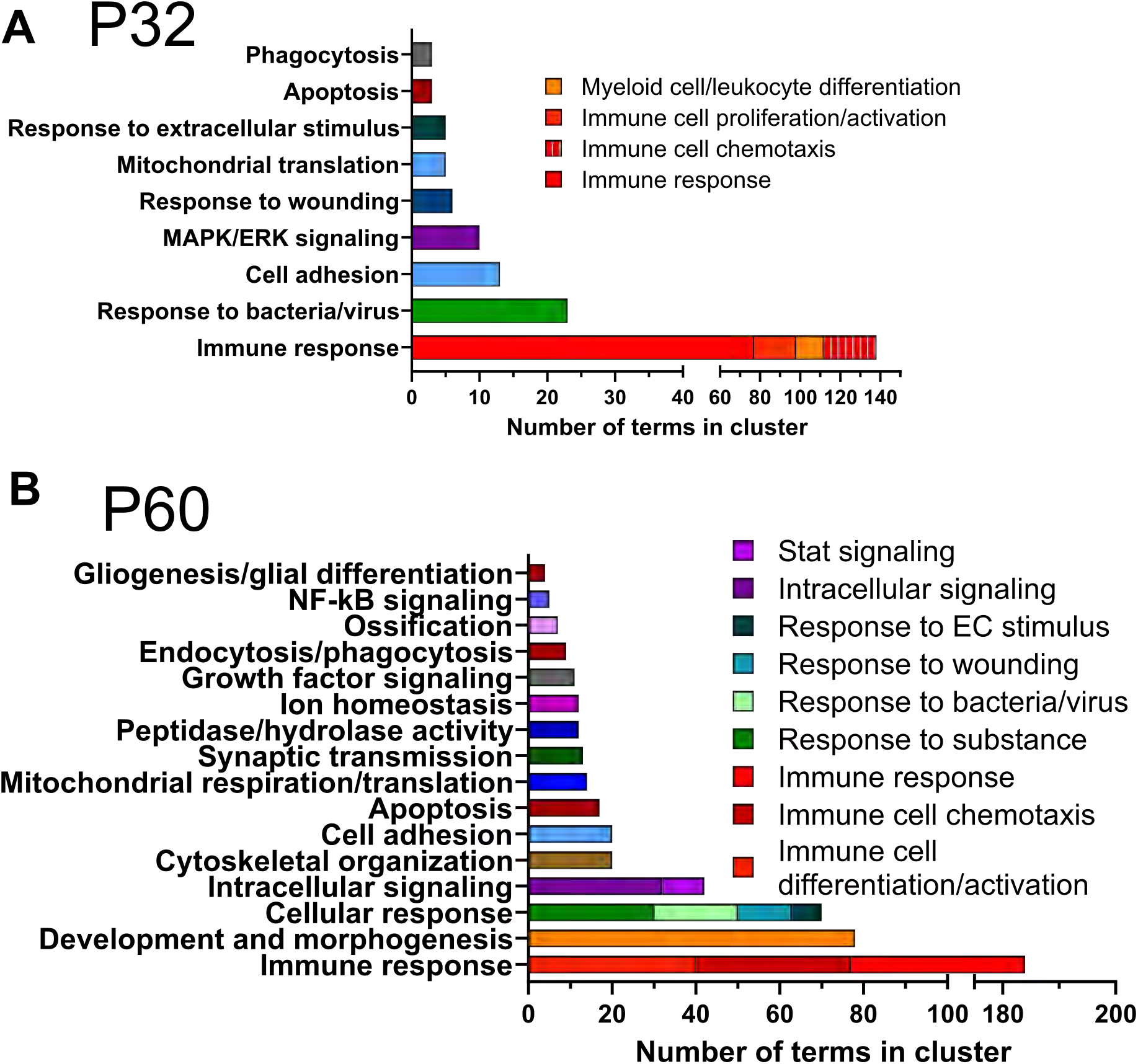
Number GO categories in each identified cluster in the GSEA networks presented in Figure 2. **A:** Number of categories in each cluster when comparing P32 hearing to P32 deaf. **B:** Number of categories in each cluster when comparing P60 hearing to P60 deaf. EC = extracellular

**Supplemental Figure 4.**
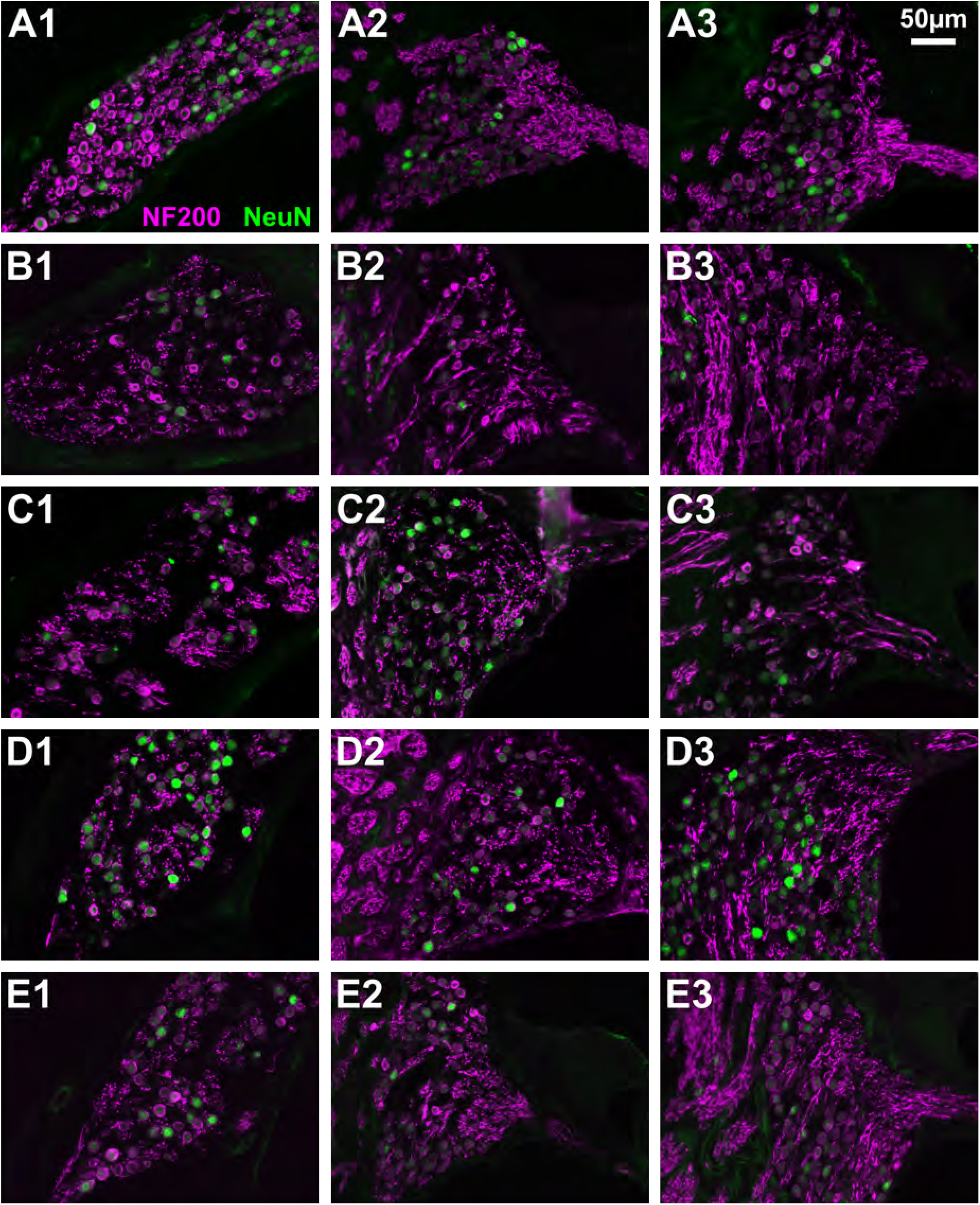
Representative images showing additional turns of the spiral ganglion in P70 hearing and deafened rats. Rats were deafened with kanamycin and received daily injections of vehicle, P7C3, ibuprofen, or P7C3+ibuprofen as described in Materials and Methods. Rats were killed and 6µm thick cryosections prepared and labeled with anti-NF200 (magenta) and anti-NeuN (green). Images show the basal (column 1), middle 2 (column 2), apical (column 3) turns from hearing **(A)** and deafened rats treated with vehicle **(B)**, P7C3 **(C)**, ibuprofen **(D)**, or P7C3+ibuprofen **(E)**. The corresponding quantitation is shown in figure 7G.

**Supplemental Figure 5.**
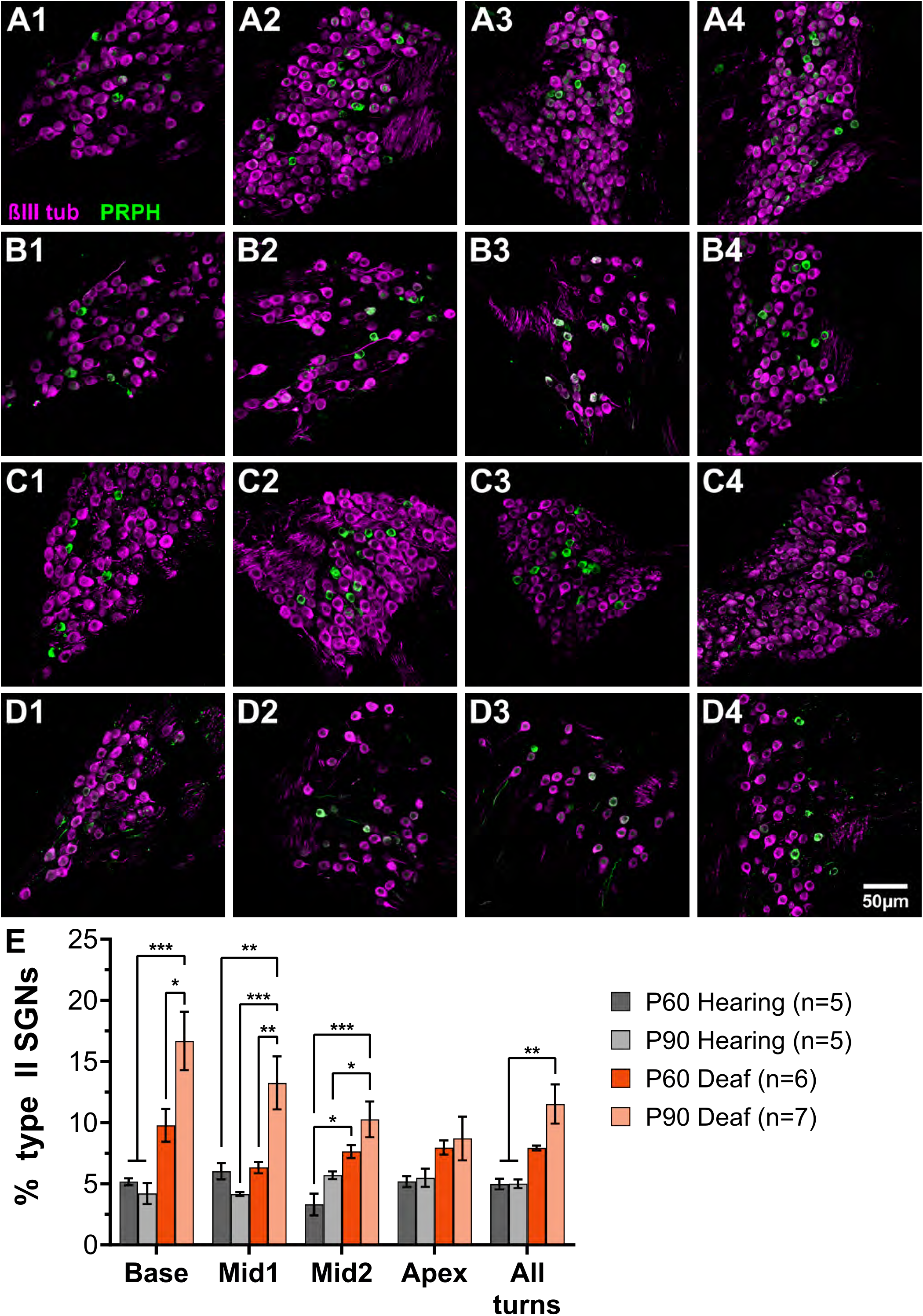
Type II SGNs are resistant to degeneration after deafening. Representative images of all SGNs (β_III_-tubulin, magenta) and type II SGNs (PRPH, green) in hearing rats at P60 **(A1-4)** or P90 **(C1-4)** and deafened rats at P60 **(B1-4)** or P90 **(D)**. Deafened rats received daily injections of kanamycin from P8-P16 as described Materials and Methods. Rats were killed at P60 or P90 and cochleae collected and cryosections prepared. The total number of surviving SGNs and the number of type II SGNs were counted and the percentage of type II SGNs for each cochlear turn was calculated. The base is shown in column 1, middle 1 in column 2, middle 2 in column 3, and apex in column 4. **E:** Quantitation of the percentage of type II SGN percentage in P60 hearing/deaf and P90 hearing/deaf. One-way ANOVA with Tukey’s multiple comparisons at each cochlear location. *p<0.05, **p<0.01, ***p<0.001; n = # of animals.

**Supplemental Figure 6.**
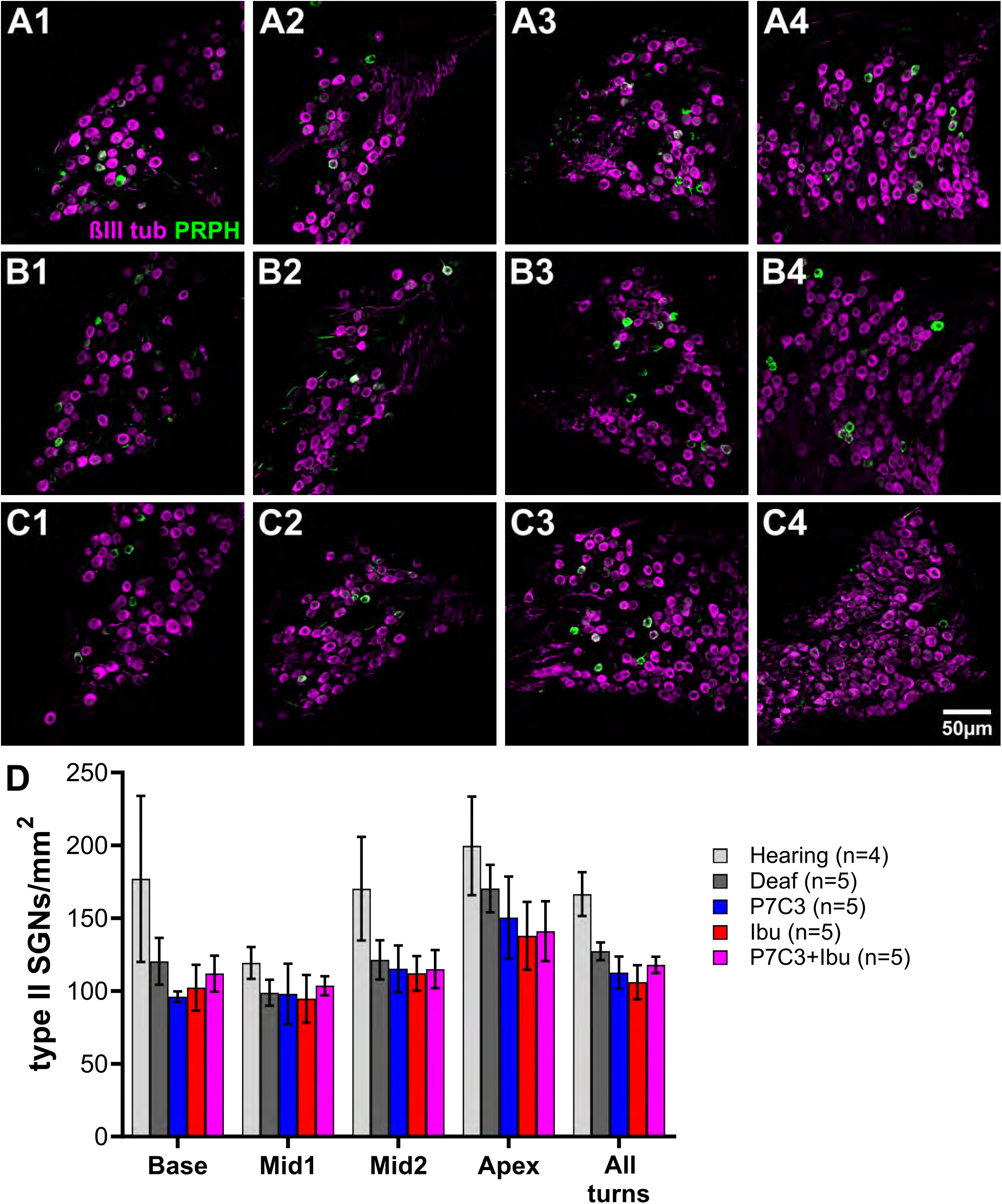
Drug treatment does not affect type II SGN degeneration. Rats were deafened with kanamycin and received daily injections of vehicle, P7C3, ibuprofen, or P7C3+ibuprofen as described in Materials and Methods. Rats were euthanized at P70 and cryosections prepared and labeled for all SGNs and type II SGNs as described in Materials and Methods. Representative images of all SGNs (β_III_-tubulin immunofluorescence, magenta) and type II SGNs (peripherin immunofluorescence, PRPH, green) in P70 deafened rats treated with P7C3 **(A1-4)**, ibuprofen **(B1-4)**, or P7C3+ibuprofen **(C1-4).** The base is shown in column 1, middle 1 in column 2, middle 2 in column 3, and apex in column 4. **D:** The graph shows quantitation of type II SGN density in all conditions and across cochlear location, n = # of animals.

